# Transcriptome and chromatin accessibility landscapes across 25 distinct human brain regions expand the susceptibility gene set for neuropsychiatric disorders

**DOI:** 10.1101/2022.09.02.506419

**Authors:** Pengfei Dong, Jaroslav Bendl, Ruth Misir, Zhiping Shao, Jonathan Edelstien, David A Davis, Vahram Haroutunian, William K. Scott, Susanne Acker, Nathan Lawless, Gabriel E. Hoffman, John F. Fullard, Panos Roussos

## Abstract

Brain region- and cell-specific transcriptomic and epigenomic molecular features are associated with heritability for neuropsychiatric traits, but a systematic view, considering cortical and subcortical regions, is lacking. Here, we provide an atlas of chromatin accessibility and gene expression in neuronal and non-neuronal nuclei across 25 distinct human cortical and subcortical brain regions from 6 neurotypical controls. We identified extensive gene expression and chromatin accessibility differences across brain regions, including variation in alternative promoter-isoform usage and enhancer-promoter interactions. Genes with distinct promoter-isoform usage across brain regions are strongly enriched for neuropsychiatric disease risk variants. Using an integrative approach, we characterized the function of the brain region-specific chromatin co-accessibility and gene-coexpression modules that are robustly associated with genetic risk for neuropsychiatric disorders. In addition, we identified a novel set of genes that is enriched for disease risk variants but is independent of cell-type specific gene expression and known susceptibility pathways. Our results provide a valuable resource for studying molecular regulation across multiple regions of the human brain and suggest a unique contribution of epigenetic modifications from subcortical areas to neuropsychiatric disorders.

## Introduction

The majority of genetic risk variants for neuropsychiatric disorders, including schizophrenia (SCZ)^1^, bipolar disorder (BD)^2^, and Autism spectrum disorder (ASD)^3^, are non-coding, and interpreting the functional consequences of these variants is challenging^4,5^. Cis-regulatory elements (CRE), including promoters and enhancers, of the central nervous system (CNS) are enriched for the non-coding risk variants^6–9^. For a given gene, promoters are regulatory units that can control the expression of associated isoforms (i.e. promoter-isoforms) independently in different contexts, such as cell types and disease status^10,11^. Enhancers are distal regulatory elements that also regulate target gene expression in a cell-type and context-dependent manner^12^. The prevailing view is that non-coding risk variants exert their effects by altering CRE function and disrupting gene-regulatory circuits^6,13^. Large-scale efforts have been fruitful in generating CRE regulatory atlases^8,9,14–16^ and mapping chromatin quantitative trait loci (QTL)^7,17,18^ in the human brain to decipher the function of neuropsychiatric risk variants.

The human brain is a highly complex organ consisting of myriad cell types that reside in specific brain regions that are associated with different cognitive functions. Accordingly, the gene expression and epigenome landscapes can vary dramatically between brain regions and cell types^9,19,20^. While previous efforts focused on the cortical areas of the brain, the molecular mechanisms regulating subcortical areas are largely unknown. Emerging evidence from genetic association^9,19,21^ and neuroimaging^22,23^ studies suggest a critical role for subcortical areas in neuropsychiatric disorders. One prominent example is basal ganglia-specific medium spiny neurons (MSN), in which both expressed genes^1,21^ and accessible chromatin^9,19^ are strongly enriched for common SCZ risk variants, and are independent of other neuronal cell populations^21^. We previously profiled chromatin accessibility across cortical and subcortical areas and highlighted the function of striatum CREs in neuropsychiatric disorders^9^. However, the previous data set included only a limited number of subcortical areas and lacked coupled gene expression information. Thus, the cell-type-specific promoter- and enhancer- regulatory landscape across brain regions, as well as the association with risk variants in neuropsychiatric disorders, remain uncharacterized.

To this end, we comprehensively profiled gene expression and chromatin accessibility in neuronal and non-neuronal nuclei across 25 brain regions from 6 individuals with no history of neuropsychiatric or neurodegenerative disease. We found extensive and consistent regional differences in gene expression and chromatin accessibility, and identified alternative promoter-isoform usage and enhancer-promoter interactions across the different brain regions. By leveraging a systems biology approach, we identified brain region-specific gene co-expression and chromatin co-accessibility modules that are robustly associated with neuropsychiatric disorders. We highlight a novel set of genes that is strongly overrepresented for both common and rare risk variants for neuropsychiatric disorders, as well as candidate causal genes identified by TWAS and statistical fine-mapping, yet is independent of reported cell type-specific gene expression and known schizophrenia-associated pathways. Lastly, we incorporate our multi-omics data with a machine-learning classifier to predict the potential functional effect of risk variants in neuropsychiatric disorders. Our data provide an important resource for understanding the regulome and molecular mechanisms in neuropsychiatric traits.

## Results

### An atlas of chromatin accessibility and gene expression in neuronal and non-neuronal nuclei across 25 brain regions

To comprehensively characterize gene expression and chromatin accessibility across cortical and subcortical structures, we generated matched ATAC-seq and RNA-seq profiles in neuronal and non-neuronal nuclei isolated from 25 brain regions covering the forebrain (ForeBr), basal ganglia (BasGan), midbrain/thalamus (MidBr) and hindbrain (HinBr) of six control subjects (**Fig. 1a,b** and **Fig. S1**). After rigorous quality control, including assessing cell type, sex, genotype concordance, and quality metrics (**Fig. S2** and **Supplementary Notes**), we obtained a total of 14.2 billion uniquely mapped paired-end read pairs for RNA-seq (N=265), and 16.1 billion uniquely mapped paired-end read pairs for ATAC-seq (N=210). We examined the abundance of neuronal brain region marker genes, including *KCNS1* (ForeBr), *DRD2* (BasGan), *IRX3* (MidBr), and *GABRA6* (HinBr), and found concordant brain region specificity for both gene expression (**Fig. S3a**) and chromatin accessibility (**Fig. 1c**). We generated an atlas of open chromatin regions (OCRs) by calling peaks across 25 brain regions for each cell type (**Fig. S3b**), yielding 320,308 and 196,467 neuronal and non-neuronal peaks, respectively. To further confirm the brain-region and cell-type specificity of OCRs, we compared our results with a recent multi-brain-region single-cell ATAC-seq dataset^19^. In line with the previous analyses^9,19^, both neuronal and non-neuronal OCRs were enriched in their corresponding cell types, and neuronal OCRs exhibited brain region specificity (**Fig. S3c**), as previously reported.

**Fig. 1.**
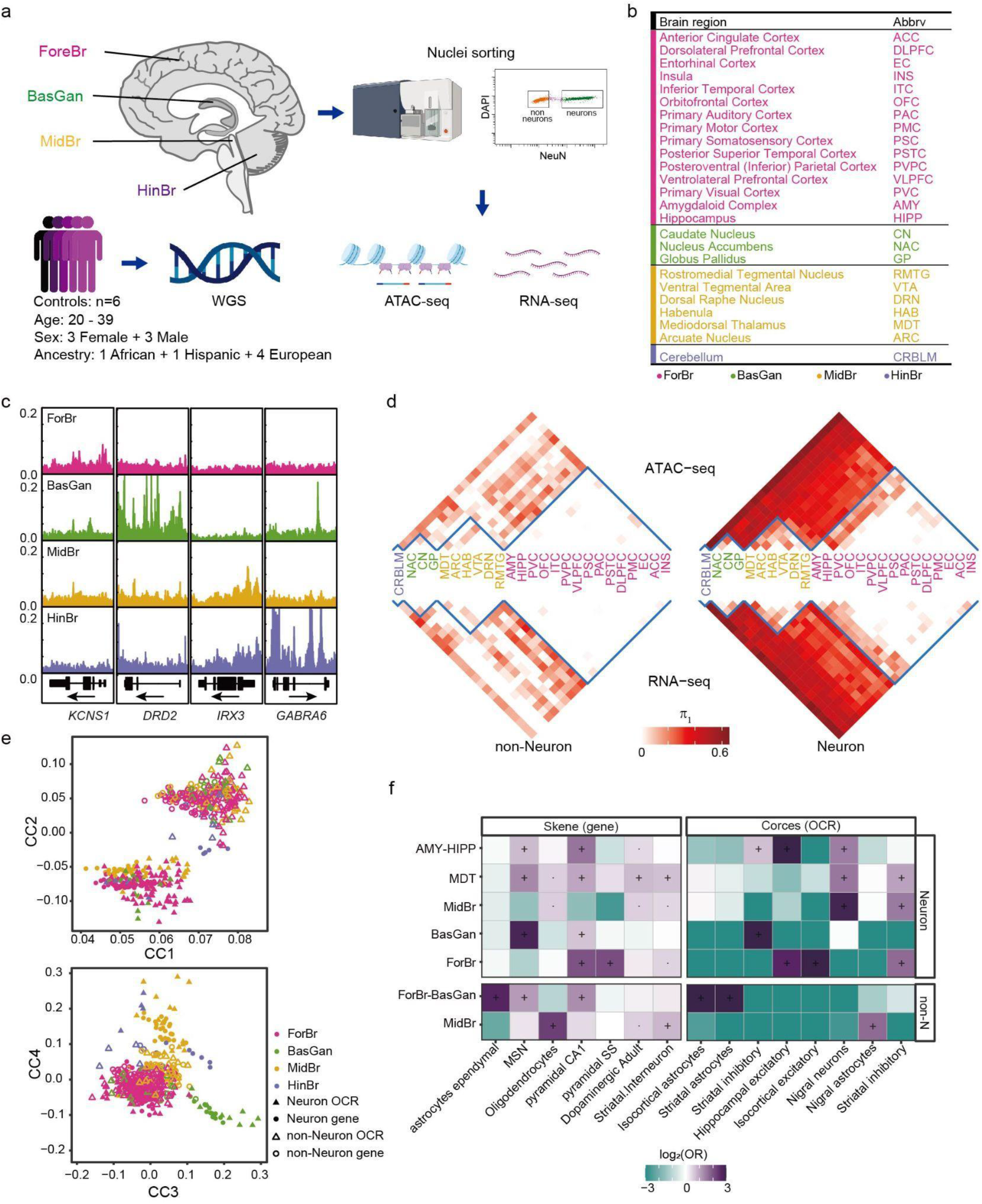
Extensive and consistent gene expression and chromatin accessibility across 25 human brain regions. **a**. Schematic representation of study design. Postmortem samples were dissected from 25 brain regions across ForeBr, MidBr, BasGan, and HindBr. Nuclei were subjected to fluorescence-activated nuclear sorting (FANS) to yield neuronal (NeuN+) and non-neuronal (NeuN-) nuclei, followed by ATAC-seq and RNA-seq profiling. We also performed whole-genome sequencing for each individual. **b**, The 25 brain regions and abbreviations (anatomy dissection in **Fig. S1**). **c**, Chromatin accessibility profiles merged from the neuronal broad brain regions around brain region-specific marker genes. **d**, Pairwise statistical dissimilarity (quantified based on the proportion of true tests, π1) across different brain regions of neuronal and non-neuronal cells in the two assays. **e**, the shared correlation structures (canonical correlation vectors, CC) between ATAC-seq and RNA-seq are separated by cell type (top, CC1-2) and brain regions (bottom, CC3-4). **f**, Cell type enrichment in brain region-specific gene (for DEG)^21^ and OCR (for DAC)^19^ sets. non-N represents non-neurons. Odds ratio (OR). “·”: Nominally significant (*P* < 0.05); “+”: significant after FDR (Benjamini & Hochberg) correction (FDR < 0.05).

### Consistent transcriptomic and chromatin accessibility differences across brain regions

To visualize the genome-wide molecular feature similarities between cell type and brain regions, we applied multidimensional scaling (MDS) analysis to both gene expression and chromatin accessibility (**Fig. S3d**). Consistent with our previous analysis, samples clustered first by cell type and next by brain region in both assays^9^. We performed differential analysis for genes and OCRs for each pair of brain regions and used Storey’s π_1_ statistic^24^ as a metric of dissimilarity across brain regions (**Fig. 1d**). In neurons, both assays exhibit robust differences across broad brain regions that include ForeBr, BasGan, MidBr, and HinBr. A subset of brain regions, including the amygdala (AMY) and hippocampus (HIPP) in the ForeBr, as well as mediodorsal thalamus (MDT) in the MidBr, exhibit substantial differences compared to other areas within the same broad brain region (**Fig. 1d**). In non-neurons, MidBr exhibit robust differences compared to the other broad brain regions (**Fig. 1d**). The HinBr exhibits the most significant differences in comparison to the other brain regions. However, given the challenge of separating neuronal from non-neuronal nuclei in HindBr^25^, we provided HindBr RNA-seq and ATAC-seq as a resource but did not include these data in downstream analyses.

To determine the relationship between the two molecular features, we assessed the genome-wide consistency between gene expression and the corresponding promoter chromatin accessibility. Consistent with previous transcriptional regulation models^26–28^, we observe a high correlation between gene expression and promoter chromatin accessibility (**Fig. S3e**). To further assess the shared correlation structure between RNA-seq and ATAC-seq, we performed canonical correlation analysis (CCA)^29^ and found the samples are separated by cell type and brain region instead of assays (**Fig. 1e**), demonstrating the consistency of both approaches.

We found that, in neurons, 87.5% (19,151/21,878) of genes and 79.6% (235,865/296,337) of OCRs are significantly differentially expressed/accessible in at least one of the pairwise comparisons between ForeBr, BasGan, and MidBr. In order to be conservative, genes/OCRs were considered broad-brain-region specific only if they were significantly more expressed/accessible in all pairwise comparisons against the remaining broad-brain-regions. Consistent with our previous analysis^9^, BasGan has the highest number of differentially expressed genes (DEGs) and differentially accessible chromatin regions (DACs) (**Fig. S4a**). Interestingly, neuronal BasGan-specific genes contain a significantly higher fraction of noncoding RNAs. One such example is the *MALAT1* gene, which has been implicated in multiple neuropsychiatric traits^30^ (**Fig. S4b**).

To further examine the brain-region and cell-type specificity of our findings, we compared our differential analysis with two independent multi-brain-region single-cell reference sets^19,21^. Consistent with known brain region anatomy, the DEGs from ForeBr, BasGan, and MidBr were enriched for pyramidal cells, MSN, and dopaminergic neurons, respectively (**Fig. 1f**). The DACs exhibit similar cell-type associations and show stronger specificity and enrichment (**Fig. 1f**). In non-neurons, the MidBr-specific DEGs and DACs were both associated with astrocytes (**Fig. 1f**), consistent with a recent report of astrocyte heterogeneity between the MidBr and other brain regions^31^. We further assessed the association between DEGs/DACs and common neuropsychiatric disorder-associated variants (**Methods**). Consistent with the previous analysis, the neuronal brain region-specific molecular features were robustly associated with multiple neuropsychiatric traits (**Fig. S4d**). As non-neurons exhibit limited brain region specificity (**Fig. 1d**) and show limited association with risk loci for neuropsychiatric traits (**Fig. S4d**), we focus only on neurons for downstream brain regional analyses.

### Alternative promoter isoform usage across brain regions identified neuropsychiatric disorder susceptible gene sets

Usage of alternative promoters regulates isoform usage pre-transcriptionally^10,11^. Promoter isoform expression can be quantified by examining the set of unique junction reads transcribed from the promoter using RNA-seq data^32^. As full-length isoforms significantly outnumber promoter-isoforms, the quantification of promoter-isoform expression is substantially more robust^32^. By modeling the junction reads that are uniquely identifiable for the first exon (**Fig. 2a** and **Methods**), we detected uniquely identifiable 13,108 promoter-isoforms (9,108 5’ promoters and 4,000 non-5’ promoters) from 11,224 genes. Although without regulatory genomics annotation, the 5’ promoter is often assumed as the default promoter, we found, at least for a fraction of genes (1,344/11,224), that the 5’ promoter-isoform is not the most highly expressed (i.e., major promoter, **Fig. S5a**). To validate our annotation of the non-5’ major promoters, we utilized independent promoter-associated epigenomic data (including ATAC-Seq, and H3K4me3 and H3K27ac ChIP-Seq), around both the 5’ promoter and the non-5’ major promoter, of relevant genes in neurons. As expected, the major promoters (non-5’) exhibit more potent active epigenomic modifications than the 5’ promoters (**Fig. S5b**), validating our promoter-isoform result.

**Fig. 2.**
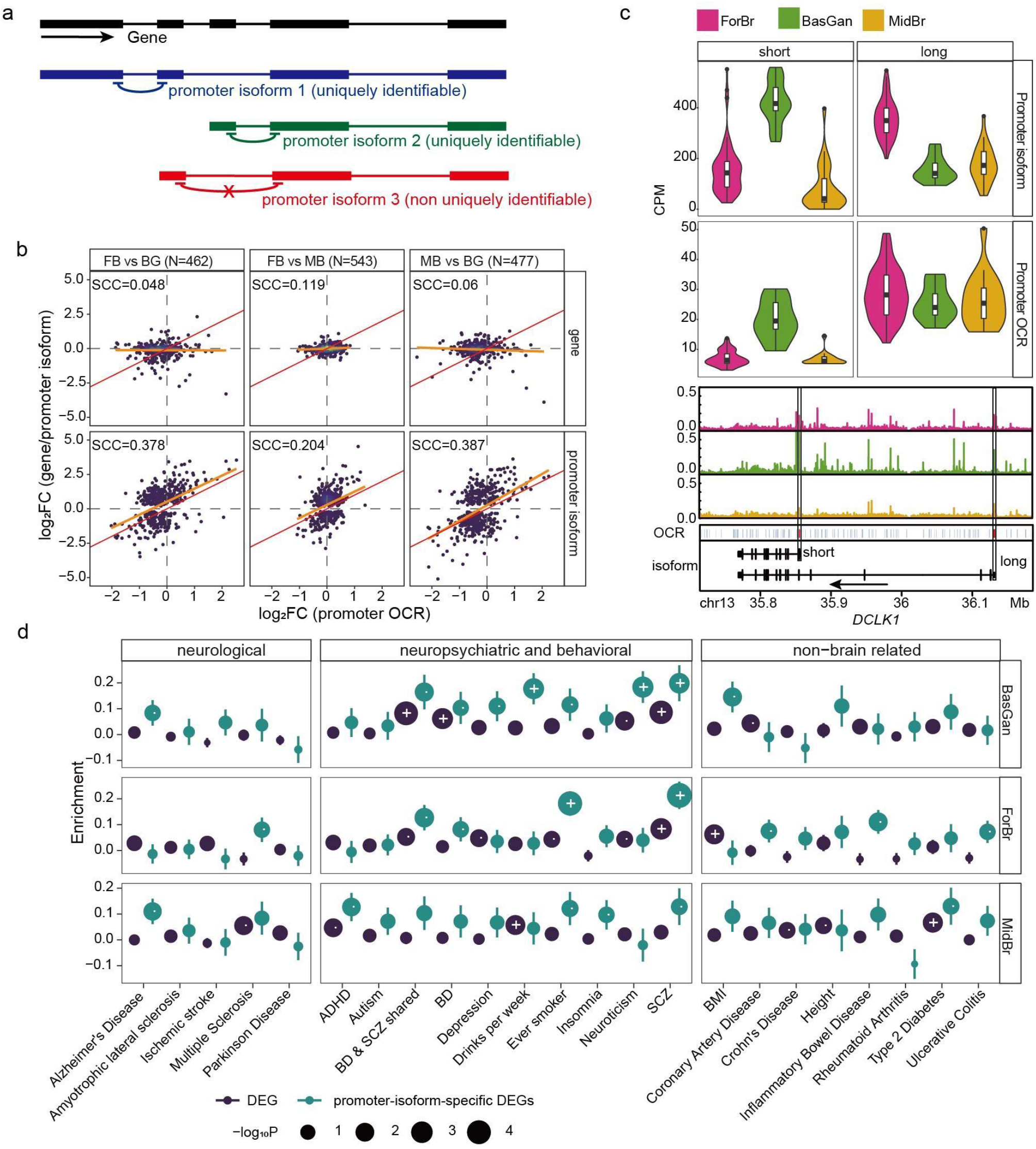
Promoter isoform alternations between brain regions. **a**, Schematic representation of promoter-isoform quantification using RNA-seq data from neurons. Transcripts that are regulated by the same promoter are grouped (promoter-isoform), and are quantified using the set of unique junction reads spanning the first intron of each transcript. The non-uniquely identifiable promoter-isoforms (red) were removed from this analysis (**Methods**). **b**, Spearman correlation coefficient (SCC) between the log2 fold change of gene/promoter isoform and promoter OCR across brain regions. N indicates the number of non-concordant promoter-isoforms. **c**, upper panel, the promoter isoform expression and promoter OCR chromatin accessibility level across brain regions. Bottom panel, chromatin accessibility profiles (neuronal) and the position of the two promoter-isoforms, and promoter OCRs (highlighted in red bars and black boxes). **d**, enrichment of common variants for different trait classes at the DEGs, and promoter-isoform-specific DEGs for each brain region. The y-axis represents the enrichment level (MAGMA beta ± standard error), and the size represents the *P* value. “·”: Nominally significant (p<0.05); “+”: significant after FDR (Benjamini & Hochberg) correction (FDR < 0.05).

To examine promoter-isoform-specific brain region alterations, we compared neuronal DEGs between promoter-isoform and the parent gene across brain regions. As expected, the majority of promoter-isoforms are concordant with the parent genes (**Fig. S5c**). A fraction of promoter-isoforms is significantly differentially expressed between brain regions (FDR < 0.05), while the parent gene is either not significant (nominal *P* > 0.1), or shows the opposite direction (promoter-isoform specific DEG, **Fig. S5c** and **Methods**), suggesting an alternative promoter-isoform usage across brain regions. In contrast to the majority of genes (**Fig. S3e**), the parent gene expression of these promoter-isoforms exhibits a very limited association with promoter OCRs (**Fig. 2b; top panel**); while promoter-isoform expression correlates well with promoter OCRs across brain regions (**Fig. 2b; bottom panel**), providing additional support for using promoter-isoform, rather than gene expression, to explore transcriptome differences across brain regions. One prominent example is the Doublecortin Like Kinase 1 (*DLK1*) gene, an essential regulator of synaptic development and axon regeneration that consist of two promoter-isoforms (a long and short form, respectively, **Fig. 2c**)^33^. The protein corresponding to the short isoform acts as an inhibitor of the long isoform protein, and this inhibition is regulated by calcium concentration^34^. The long isoform is highly expressed in ForeBr and has the lowest expression in BasGan. In contrast, the short isoform is highly expressed in BasGan. Consistent with this, the promoter OCR of the short isoform also has higher chromatin accessibility in BasGan (**Fig. 2c**).

More importantly, compared to brain region-specific DEGs, promoter-isoform-specific DEGs have a markedly higher enrichment for common risk variants for multiple neuropsychiatric traits, including SCZ (**Fig. 2d**). Moreover, the promoter-isoform-specific DEGs are strongly enriched for rare coding variants for SCZ and ASD (**Fig. S5d**). Consistent with this, such genes are strongly overrepresented for synaptic functions (**Fig. S5d**). Our analysis highlights the alternative promoter usage across different brain regions and their role in neuropsychiatric disorders.

### Enhancer-promoter links explain variation in gene expression

Most of the OCRs in the human brain are outside of the promoter areas (**Fig. S3b**), and are strongly enriched for neuropsychiatric and neurological disorder risk variants^8^. As the mammalian genome is organized in three-dimensional (3D) space, and promoter-enhancer (EP) links can span considerable genomic distances^12^, determining enhancer-gene regulation is critical for understanding the molecular mechanisms of complex traits. The recently developed activity-by-contact (ABC) model integrating chromatin states and 3D contacts accurately predicts the experimentally validated promoter-enhancer interactions^35^. With our brain-region-specific chromatin accessibility information and cell-type-specific Hi-C contacts^18,36^, we built brain-region-specific ABC models and found the majority of EP links located within 50kb of transcriptional start sites (TSS) (**Fig. S6a** and **Supplementary Table 8**). Consistent with gene expression and chromatin accessibility, EP interactions also exhibit considerable regional-specificity in neuronal cells across brain regions (**Fig. 3a**). As the model only considered physical proximity and epigenomic signal strength, we next assessed the coordination between EP links. We compared the Pearson correlation between gene expression and chromatin accessibility of brain region-specific EPs, shared EPs, and non-EPs (**Fig. 3b**). For each of the brain regions, brain region-specific-EPs exhibit similar correlations compared to shared EPs, while the background (other brain region-specific, non-EP) is much lower, validating our brain region-specific EP links.

**Fig. 3.**
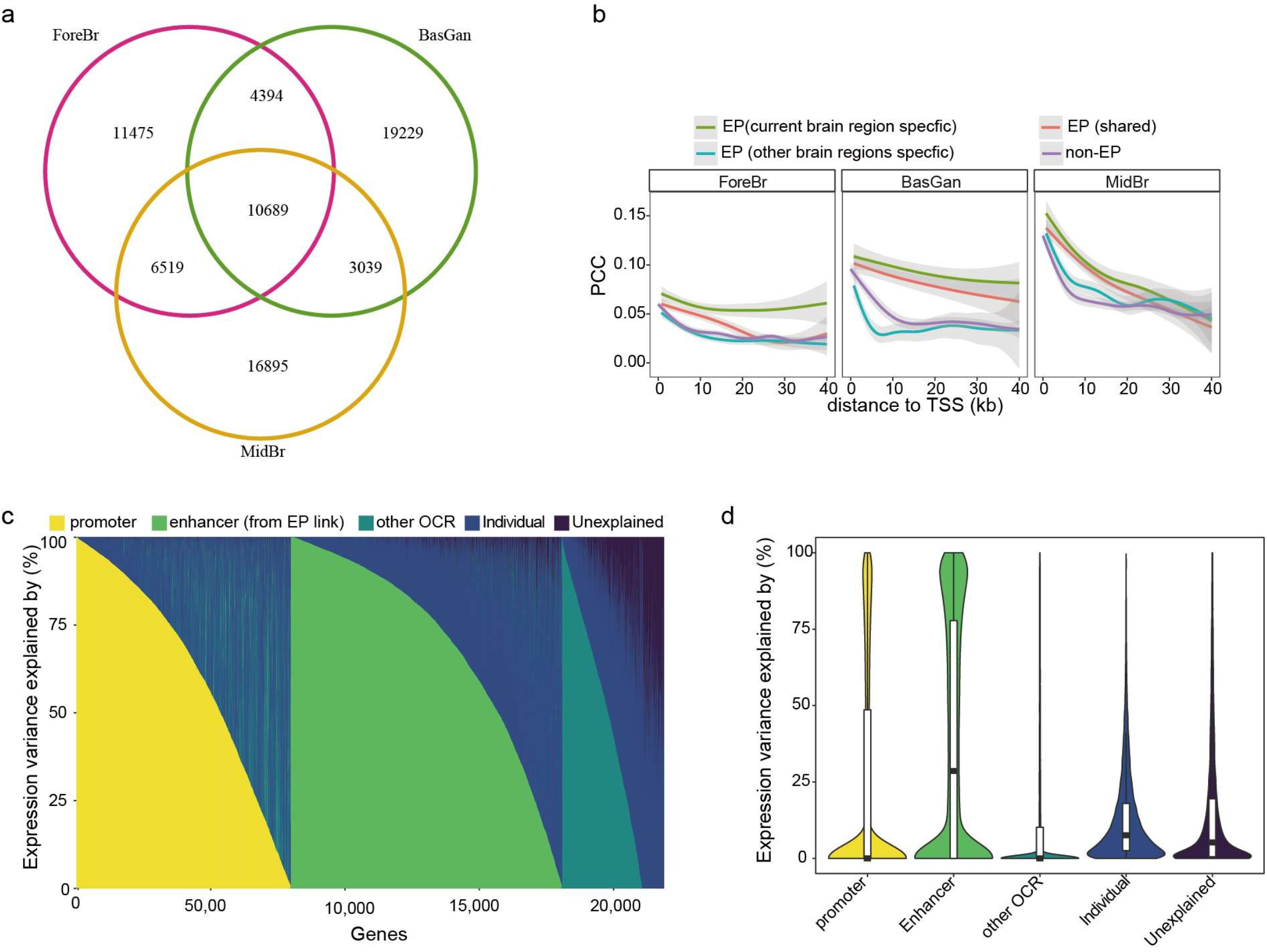
Brain region-specific neuronal gene-enhancer links. **a**, Overlap of ABC gene-enhancer links among brain regions. **b**, Correlations between gene expression and enhancer activity of shared ABC, brain region-specific ABC, other brain region-specific ABC, and non-ABC pairs for the three brain regions. The line shows the mean, and the shadow shows the 95% confidence intervals. **c**, Variance component decomposition of gene expression (every gene per column) by the promoter, ABC-enhancer (ABC), other OCRs within 200kb (other OCR), and Individual, and **d**, the associated variance distribution across different categories.

To further examine the contribution of different types of CRE (i.e. promoters, EP-linked-enhancers, and other-OCRs) to gene expression, we fit a variance component mode^l37,38^ for every gene to quantify the proportion of expression variation explained by components of the genome-wide chromatin accessibility landscape. Although EP-linked-enhancers only account for a small fraction of OCRs (12.7% i.e. 37,817/296,337), they explained the highest proportion of gene expression variance (**Fig. 3c,d** and **Fig. S6b**). Together with promoters, more than half of the gene expression variance could be explained. Interestingly, the EP-enhancer-explained brain-region-specific genes are more strongly enriched for CNS cell types and synaptic pathways compared to those of the promoter-explained, and other factors explained, demonstrating the importance of enhancer regulation in cell-type-specific functions in the human brain (**Fig. S6c**).

### Co-accessibility network refines neuropsychiatric susceptible genes complementary to gene co-expression analysis

As neuropsychiatric traits are highly polygenic, their etiology is organized into functional groups and is expected to have a shared genetic component^39,40^. In addition to gene co-expression networks, chromatin co-accessibility or CRE co-activity analysis has emerged as a means to provide new insights into regulatory mechanisms in development^41,42^ and disease^43^. As such, we constructed weighted gene co-expression analysis (WGCNA) networks with genes, promoters, and enhancers (EP-linked enhancers), to systematically examine the co-expression and co-accessibility modules implicated in disease etiology (**Fig. S7a** and **Methods**). Together, we identified 19 gene modules, 7 promoter modules, and 11 enhancer modules. We found that 89.5% (17/19) of the gene modules, and all of the gene/enhancer modules, are preserved in independent data sets (**Fig. S7b** and **Methods**). Although constructed independently, the gene, promoter, and enhancer modules captured consistent molecular variation across human brain regions, as demonstrated by unsupervised hierarchical clustering (**Fig. 4a**). As expected, the linked genes (for promoter and enhancer modules) and linked CREs (for gene modules) exhibit concordant brain-region-specificity enrichment. However, the co-expression module genes and the co-accessibility module-linked genes are not entirely overlapping (**Fig. S7c**), suggesting that chromatin co-accessibility analysis will provide new information to complement the co-expression network.

**Fig. 4.**
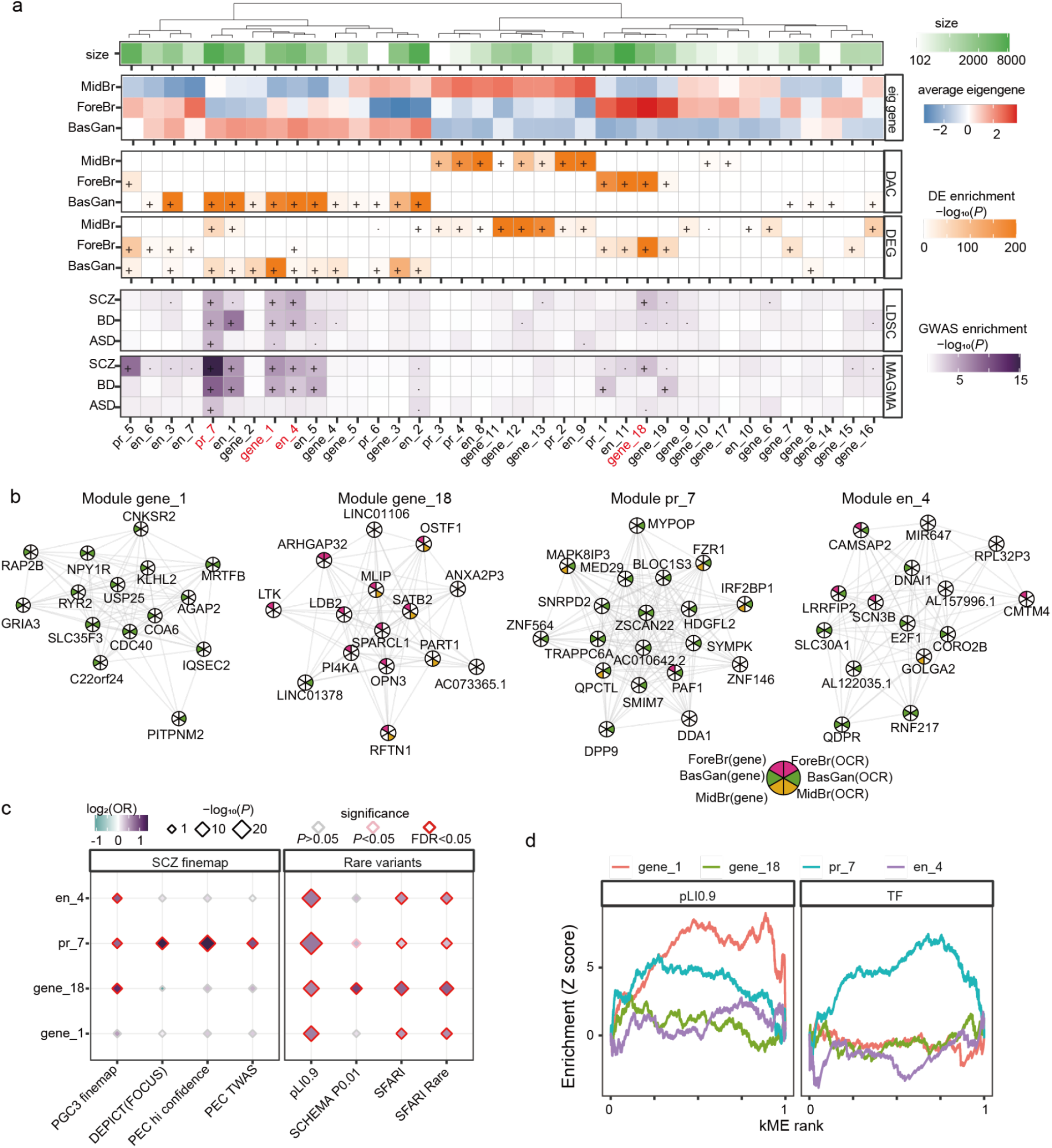
WGCNA network. **a**, Unsupervised hierarchical clustering of gene, promoter, and enhancer modules by the eigengene. Brain-region (DEG, DAC), and neuropsychiatric disorder common variants (ASD, BD, and SCZ) enrichment across gene/CRE modules as well as the associated CRE/genes. LDSC and MAGMA were used to determine the GWAS enrichment in CRE and genes, respectively. “·”: Nominally significant (p<0.05); “+”: significant after FDR (Benjamini & Hochberg) correction (FDR<0.05). **b**, the top 25 hub genes/CRE (linked genes shown) are shown for the prioritized modules. Edges represent coexpression. Nodes are colored to represent brain-region specific gene expression, SCZ implicated pathways genes, and TF genes. **c**, rare variants, loss of function intolerant genes(pLI0.9), and SCZ fine-mapped gene enrichment for each of the modules. **d**, Gene-set enrichment analysis plots for LoF-intolerant genes, SCZ pathway implicated genes, and TF genes for the four modules. The extent of deviation from the horizontal reflects the degree of enrichment (+) or depletion (−) at a given centrality (kME, i.e. module memberships).

To prioritize the modules that are associated with risk variants for neuropsychiatric traits, we utilized two independent methods (LD score regression^44^ and MAGMA^45^) to assess the enrichment of modules with three important neuropsychiatric disorders: ASD^3^, BD^2^, and SCZ^1^. The disease association between module members (gene/CRE) and linked molecular features (CRE/gene) are highly consistent between both methods (**Fig. 4a**). We prioritized four modules that are robustly associated with common disease variants, including three BasGan modules (gene_1, pr_7, and en_4), and a ForeBr module (gene_18). Notably, the OCRs within pr_7 and en_4 are strongly overrepresented for disease risk variants; for example, pr_7 and en_4 exhibit >50 times per SNP heritability than the baseline model for ASD and SCZ, respectively (**Fig. S7d**). In addition, BasGan module pr_7 is strongly associated with all three disorders (**Fig. 4a** and **Fig. S7d**), suggesting a consistent molecular mechanism.

The most highly connected genes (i.e. hub genes) within modules gene_1 and gene_18 are brain-region-specific genes that are associated with synaptic functions and are implicated in neuropsychiatric disorders. The BasGan module gene_1 hub genes include neurotransmitter receptor genes *GRIA3* and *NPY1R*, where *GRIA3* is significantly associated with both rare and common variants of SCZ^1,46^; The ForeBr module gene_18 hub genes include neuron-associated GTPase-activating protein (*ARHGAP32*) and neuron development regulator *SATB2*. However, the associated promoter OCRs for gene_18 hub genes do not exhibit concordant brain-region specificity (**Fig. 4b**). Similarly, the hub CRE-associated genes (referred to as hub genes) from pr_7 and en_4 are not concordant with the promoters/enhancers. The en_4 hub genes are associated with different aspects of functions implicated in neuropsychiatric disorders, including ion channels (*SCN3B*, *SLC30A1*), and neuron development (*CAMSAP2*, *DNAI1*). However, multiple pr_7 hub genes are associated with regulatory functions, including transcription factors (TF; *ZNF564*, *ZSCAN22*, and *ZNF146*), RNA polymerase II subunit (*PAF6*), mediator (*MED29*), and the polyadenylation regulator (*SYMPK*), contrasting the synaptic genes in module gene_1, gene_18, and en_4.

Having shown that the four modules are enriched for common risk variants for neuropsychiatric disorders, we further examined the enrichment of potentially causal genes (genes nominated by statistical fine-mapping and TWAS) and rare coding variants. Although the four fine-mapped gene sets, including Psychiatric Genomics Consortium (PGC3 finemap)^1^, DevEloPing Cortex Transcriptome(DEPICT(FOCUS))^47^ and PsychENCODE (PEC hi confidence^7^, and PEC TWAS^48^), were analyzed with ForeBr or fetal tissues^1,7,47,48^, the BasGan module pr_7 genes are overrepresented in all the four gene sets (**Fig. 4c**). Moreover, the pr_7 genes are also enriched for rare ASD and SCZ^46^ associated mutations (**Fig. 4c**). Module gene_18 exhibits the strongest association for the gene sets of the fine-mapped common variants^1^ and rare-variants^46^ from the most recent large-scale analysis. In addition, all four modules were overrepresented for genes that are intolerant to loss-of-function mutations (defined as pLI > 0.9)^49^ (**Fig. 4c**), demonstrating the modules are under a higher level of purifying selection than average. We further examined the relationship between the enrichment of loss-of-function mutations and module membership and found that BasGan modules gene_1 and pr_7 are strongly enriched for LoF intolerant genes in the network center (**Fig. 4d**). As pr_7 hub genes are associated with regulatory function, we also examined the enrichment of TF genes (from Gene ontology annotation, see **Methods**), as one of the prominent categories of regulatory genes, and found that only pr_7 exhibits a significant association (**Fig. 4d**). Together, the identified three BasGan (gene_1, pr_7, and en_4) and gene_18 ForeBr modules are broadly enriched for common variants, and rare variants for neuropsychiatric disorders, as well as LoF intolerant genes.

### Module pr_7 is independent of known pathways and cell types

To examine the functional relationship between the selected modules and neuropsychiatric disorders, we determined the enrichment of common SCZ variants for the top enriched pathways of the selected modules (**Fig. 5a**). Module gene_1, gene_18, and en_4 were associated with synaptic functions that are enriched for SCZ common variations (**Fig. 5a**). The BasGan module gene_1 was enriched for postsynaptic function, ForeBr module gene_18 was associated with neurodevelopment, and en_4 linked genes were strongly enriched for different synaptic functions, consistent with their hub genes. To further dissect the SCZ-associated biological pathways, we determined the pairwise overlap between the top 50 biological pathways that are enriched for SCZ common variants and identified four broad categories of pathway groups (**Fig. 5b** and **Methods**), including calcium channel, cation channel, postsynapse, and neuron development. The enhancer module en_4 is broadly enriched in all four groups, suggesting the importance of enhancer regulation in neuropsychiatric disorders. In contrast to the postsynapse group, which was broadly enriched for all the selected modules, the other three groups exhibit module specificity. The neuron development group is strongly enriched for ForeBr module gene_18, but is not enriched for BasGan module gene_1, consistent with the analysis above describing ForeBr, but not BasGan, DEGs enrichment for neuron development (**Fig. S4c**). Our analysis suggests different brain regions might contribute to independent pathways for neuropsychiatric disorders.

**Fig. 5.**
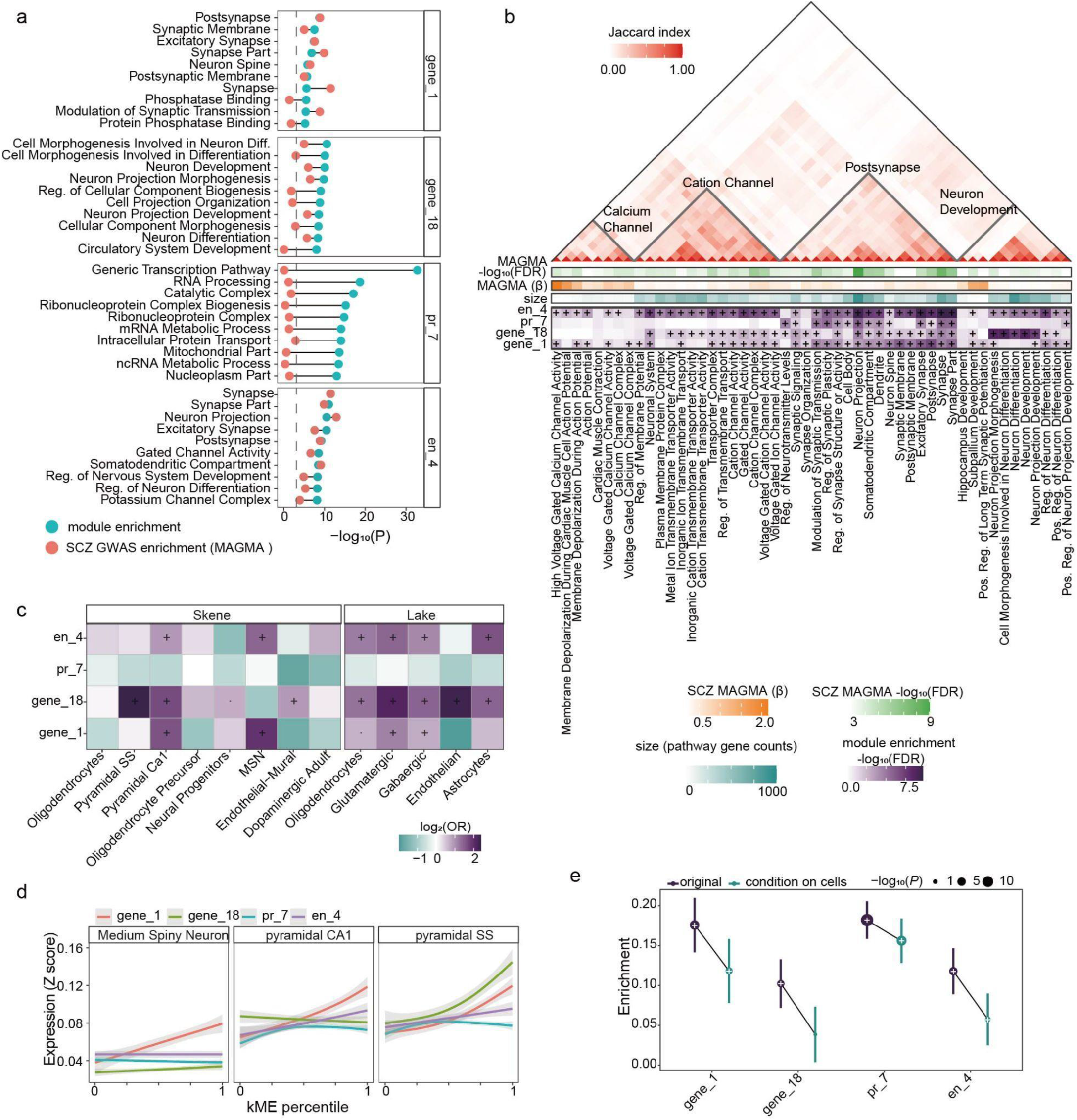
pathway and cell type of the selected modules. **a**, Top 10 enriched pathways for each prioritized module, and the pathway enrichment in SCZ common variants (MAGMA, significant after FDR Benjamini & Hochberg correction, FDR<0.05). **b**, top panel. The pairwise Jaccard index between the top 50 pathways enriched for SCZ common variants (MAGMA). The grey triangle represents the four pathway clusters identified by hierarchical clustering. Bottom panel. MAGMA enrichment (-log10(FDR), Benjamini & Hochberg correction), MAGMA enrichment coefficient(β), number of genes(size), and module enrichment for the top 50 pathways. **c**, cell type enrichment across the module with single-cell reference from humans (Lake)^50^ and mice (Skene)^21^. **d**, relative expression (Z-score, mean ± standard error, loess regression) level for most enriched cell types as a function of module membership (kME). **e**, SCZ common variants enrichment of original and condition on the expression of the reported cell types^21^ (**Methods**) across the selected modules.

We further examined the cell-type enrichment for the four modules with two single-cell references from humans^50^ and mice^21^. As expected, module gene_1, gene_18, and en_4 were enriched for critical neuronal cell-types implicated in SCZ^21^, including MSN and pyramidal neurons (**Fig. 5c**). BasGan gene_1 was most enriched in MSN cells, and ForeBr gene_18 was most enriched in neocortical somatosensory pyramidal cells (pyramidal SS), consistent with the corresponding brain regions. We further compared the cell-type-specificity^21^ with module membership and found that, except for pr_7, the remaining three modules exhibit positive associations between module membership and single-cell expression specificity (**Fig. 5d**).

In contrast to the other three modules, the top enriched pathways for pr_7 (**Fig. 5a**, and **Fig. S8c**) were related to regulatory functions, including the RNA processing, mitochondria, and catalytic function, none of which are significantly associated with SCZ common variants. In addition, pr_7 genes were depleted of marker genes from different CNS cell types (**Fig. 5c**) and were not associated with the reported cell types for SCZ (**Fig. 5d**). As such, we reasoned that the association between pr_7 and SCZ common variants is independent of the gene expression of such cell types. We thus determined the association between the genes of the four modules conditional on the expression of the reported cell types following a previous analysis^21^, including hippocampal CA1 pyramidal cells, striatal MSN, pyramidal SS, and cortical interneurons^1,21^. We found pr_7 remains highly significant and exhibits a limited decrease in enrichment following conditional analysis based on cell types (**Fig. 5e**), demonstrating that the association between pr_7 and SCZ common variants is independent of the cell-type-specific expression.

### Module pr_7 associated SCZ variants complement known pathways

To further characterize the disease variants association of module pr_7, we compared pr_7 protein-coding genes with all the significant biological pathways enriched for SCZ common variants (SCZ_pathways) (**Fig. 6a**, **Methods**, and **Supplementary Table 9**). Together, the two sets covered 7 of the 10 SCHEMA genes that contain rare coding variants that confer substantial risk for SCZ^46^ (**Fig. 6b**). Interestingly, 3 of the 4 most significant SCHEMA genes were only included in pr_7, and involve diverse regulatory functions, including epigenomic modification *(SETD1A*), ubiquitin ligation (*CUL1*), and Guanine nucleotide exchange factors (*TRIO*) (**Fig. 6b**). While the genes directly associated with synaptic function, including ion transport (*CACNA1G*, *GRIN2A*, and *GRIA3*), exhibit lower significance^46^. In addition, three extra genes with regulatory function in pr_7 were also significantly associated with rare coding mutations for SCZ (FDR < 0.05)^46^ and include genes with roles in transcription (*ZNF136*), splicesome (*SRRM2*), and epigenomic modification (*ASH1L*). Consistent with our previous analysis, the pr_7-specific genes are depleted of single cell markers and synaptic pathways but are enriched for SCZ fine-mapped genes and loss-of-function mutation intolerant genes (**Fig. 6c** and **Fig. S8d**).

**Fig. 6.**
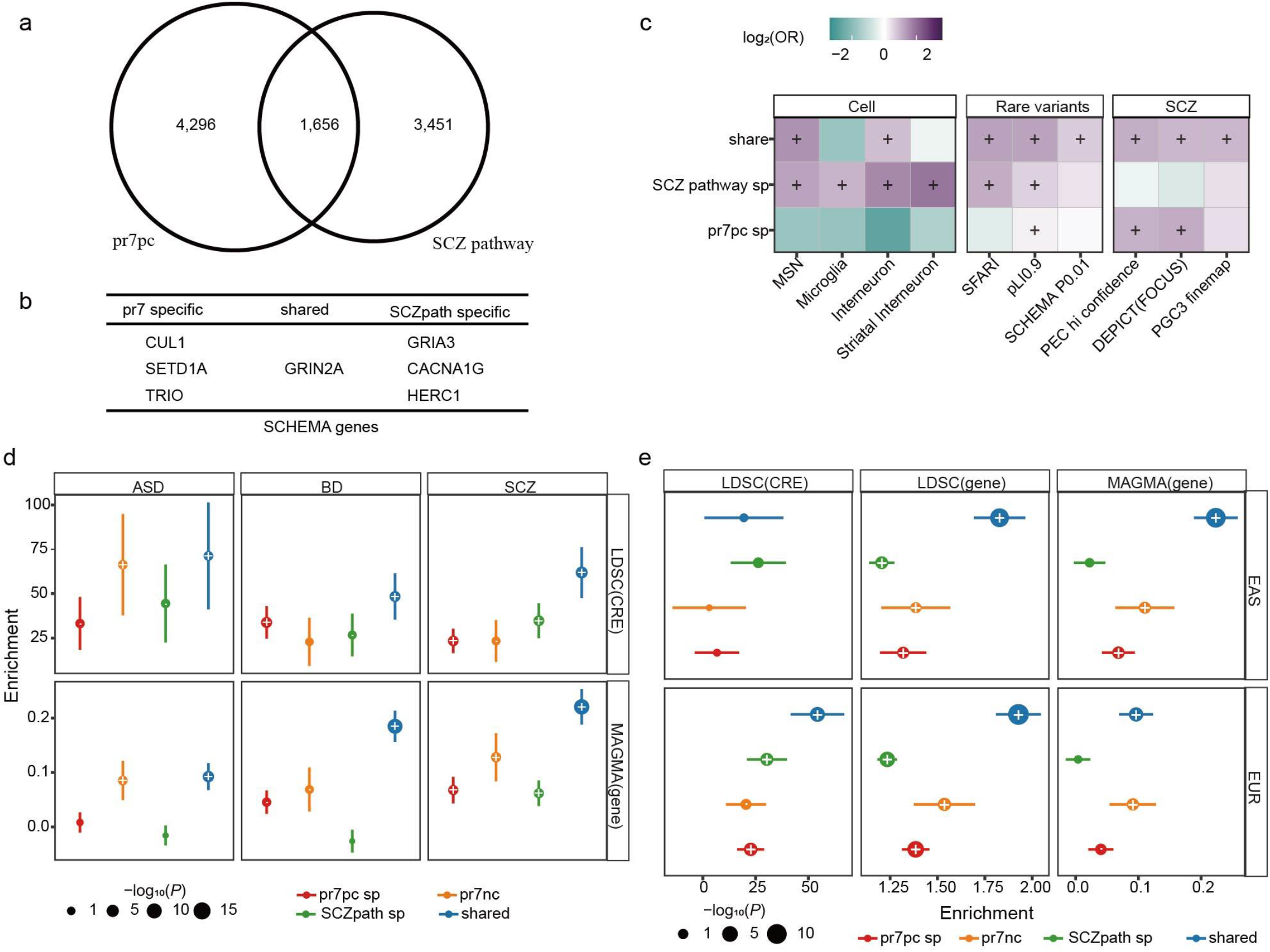
promoter module pr_7 implicated in non-canonical pathways. **a**, The Venn diagram shows the overlap between pr_7 protein-coding genes (pr7pc) and SCZ implicated pathway genes. **b**, the 7/10 SCHEMA genes implicated in the pr7pc, SCZ pathway, and shared categories. **c**, cell-type, rare variants, and SCZ fine-mapped genes for the three categories. **d**, Partitioned heritability enrichment of ASD, BD, and SCZ GWAS for different categories of genes (determined by MAGMA), and their promoter OCR (determined by LDSC). Enrichments are colored by −log_10_(*P*), and error bars represent standard error.”·”: Nominally significant (*P* < 0.05); “+”: significant after FDR (Benjamini & Hochberg) correction (FDR < 0.05). **e**, Partitioned heritability enrichment of SCZ GWAS in East Asian (EAS) and European (EUR) ancestries for different categories of genes (determined by MAGMA and LDSC) and their promoter OCR (determined by LDSC). Enrichments are colored by −log_10_(*P*), and error bars represent standard error.”·”: Nominally significant (*P* < 0.05); “+”: significant after FDR (Benjamini & Hochberg) correction (FDR < 0.05).

**Fig. 7.**
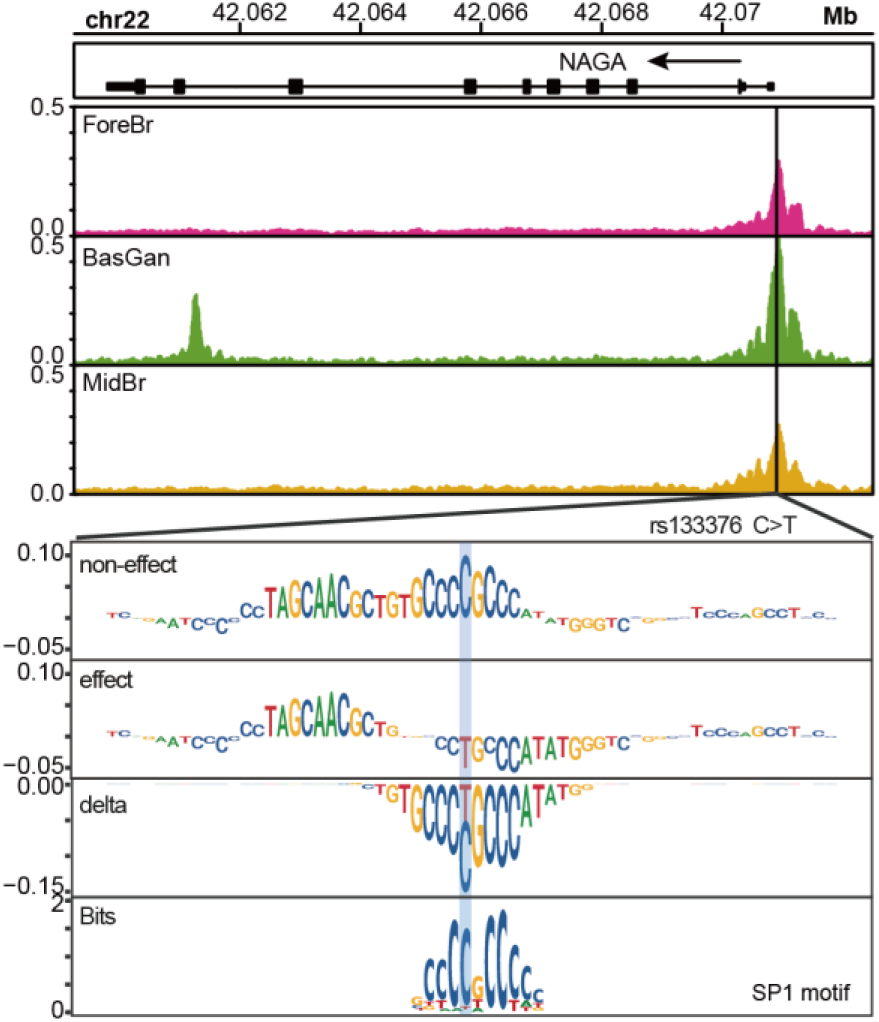
Machine learning predicts functional variants in SCZ. upper panel. ATAC-seq track around SCZ GWAS significant SNP rs133376. Bottom panel. GkmExplain importance scores for each base in the 50-bp region surrounding the SNP for the effect and non-effect alleles from the GKM-SVM model (BasGan ATAC-seq as input).

The genes shared between pr_7 and SCZ pathways are strongly enriched for the common variants of all three disorders (ASD, BD, and SCZ) (**Fig. 6c**). The pr_7-specific genes exhibit a similar level of heritability compared to the SCZ pathway-specific gene sets, suggesting that pr_7 provides an independent disease susceptibility gene set with a similar heritability level and size compared to all annotated pathways (**Fig. 6d**). In addition to protein-coding genes, pr_7 also includes 1,294 noncoding RNAs (ncRNA), including *DGCR5*, *NEAT1*, *MALAT1*, *MEG3*, and *HAR1A* that have previously been implicated in the neuropsychiatric disorders^51^. Interestingly, such ncRNAs were also significantly overrepresented for the neuropsychiatric common variants (**Fig. 6d**). To examine if the enrichment is conserved between different populations, we determined the enrichment of SCZ GWAS variants from East Asian (EAS) and European (EUR) populations and found the pr7-specific genes exhibit consistent enrichment across ancestries (**Fig. 6e**).

### Impact of regulatory variants in SCZ

As the brain-region-specific chromatin accessibility was strongly associated with neuropsychiatric traits risk variants and identified novel neuropsychiatric susceptible gene sets, we next annotated the functional effects of common variants associated with the regulatory elements identified here. We collected the SNPs that are genome-wide significant (*P* < 5 × 10^−8^) or fine-mapped causal variants (posterior probability > 0.01)^1^, as well as any SNPs in linkage disequilibrium (LD) with the 2 categories (defined as LD R^2^ > 0.8). To annotate the effect of the SNPs, we utilized a machine learning framework to score the allelic effect of an SNP on chromatin accessibility. We combined gkm-SVM^52^ with three complementary approaches, GkmExplain^53^, *in silico* mutagenesis^54^, and deltaSVM^55^, to determine the effect on chromatin accessibility in neurons across the three broad brain regions. Together, we identified 35 SNPs from 35 OCRs, including 6 promoters and 7 EP-linked enhancers that have a significant regulatory effect with all three methods (**Supplementary Table 14**). Interestingly, all 6 promoters are within the pr_7 module, and 6 of the 9 associated genes are not shared with SCZ susceptible pathways (**Supplementary Table 14**). One such example is the promoter of *NAGA*, which encodes a lysosomal enzyme that is involved in the regulation of dendritic spine pathology of SCZ^56^. The SNP is predicted to disrupt the binding motif of SP family proteins. It’s worth noting that one of the SP family TF genes *SP4* is significantly associated with both rare coding and common variants of SCZ^1,46^.

## Discussion

Understanding the brain region and cell-type-specific regulome is critical for deciphering the molecular mechanism of neuropsychiatric disorder. In this study, we present a comprehensive resource of gene expression and chromatin accessibility profiling with neurons and non-neurons across 25 brain regions in the postmortem human brain. In the neurons, we identified extensive gene expression and chromatin accessibility alterations across brain regions, which are largely driven by the distinct distribution of brain-region-specific neuronal cell subpopulations (**Fig. 1f**). The brain region-specific molecular features are strongly enriched for neuropsychiatric common risk variants (**Fig. S3d**), highlighting the brain-region-specific function in neuropsychiatric etiology. In addition to the brain-region difference in neuronal cells, we also observed consistent gene expression and chromatin accessibility alterations across brain regions in non-neuronal cells (**Fig. 1d** and **Fig. S3a**). However, as the non-neuronal brain-region-specific DEG and DAC are strongly enriched for brain-region-specific astrocytes cells from single-cell reference (**Fig. 1f**), which only constitute a limited fraction of the non-neuronal cell populations, the power of analysis to determine the heterogeneity might be restricted.

Having shown the gene expression and chromatin accessibility differences across brain regions, we reasoned that *cis*-regulation is also brain region-specific. We examined the promoter-isoform across brain regions and found although the majority of promoter-isoforms are consistent with gene expression, a substantial fraction of them represent novel differential expression patterns (i.e, only significant in promoter-isoform, or have a different sign of effect size). The promoter-isoform-specific changes were validated by the associated promoter chromatin accessibility (**Fig. 2b**), and suggest the parent genes exhibit alternative promoter usage across brain regions. Interestingly, compared to the DEGs, the promoter-isoform specific DEGs have a higher enrichment for the common variants of multiple important neuropsychiatric traits such as SCZ (**Fig. 2c**), highlighting the functional role of alternative promoter usage in neuropsychiatric traits. And such genes were strongly associated with synaptic functions. To determine the brain-region-specific enhancer regulation, we built EP links for each brain region, integrating chromatin accessibility with chromatin spatial organization. Although the linked enhancers only account for a limited fraction of total OCRs, they explained the majority of gene expression variations (**Fig. 3c**,**d**). Moreover, the enhancer-explained genes are strongly enriched for cell-type-specific marker genes.

Neuropsychiatric traits are genetically complex, and the risk variants affecting molecular features are thought to be organized into functional groups. Gene co-expression network analysis provides a powerful tool to investigate such functional groups. We identified a BasGan module (gene_1) and a ForeBr module (gene_18) that are robustly associated with SCZ common variants and ASD rare variants (**Fig. 4**). In addition, gene_18 is strongly associated with the gene sets of the fine-mapped common variants^1^ and rare-variants^46^ from the most recent large-scale analysis. It’s worth noting the two modules capture different aspects of molecular features associated with neuropsychiatric traits: the ForeBr module gene_18 is enriched for neurodevelopmental function and pyramidal cell marker genes, while the BasGan module gene_1 is enriched for post-synaptic function and MSN (**Fig. 4,5**), suggesting the risk variants might exert the function in a brain-region specific manner.

As the neuropsychiatric common variants directly affect the CRE and the downstream regulatory effect could be context-dependent, steady-state gene expression might not capture such effect^57^. Chromatin accessibility provides an alternative approach to tracing the functional effect of the genetic variants. We have further characterized the chromatin co-accessibility network. We found a promoter (pr_7) and an enhancer (en_4) chromatin co-accessbility module that is robustly associated with the risk variants of neuropsychiatric disorders. The en_4 OCRs exhibit a >50 times per SNP heritability than the baseline model for SCZ common risk variants, and the genes are strongly associated with different subclasses of SCZ susceptible pathways (**Fig. 5b**), demonstrating the importance of enhancer regulation in SCZ. The pr_7 is robustly associated with the common variants of ASD, BD, and SCZ (**Fig. S7d**). Interestingly, the genes within pr_7 are enriched for regulatory function and depleted of cell-type-specific genes. Conditional on the expression of the SCZ implicated cell types MSN, pyramidal CA1, pyramidal SS, and interneuron^1,21^, the pr_7 genes remain highly enriched for SCZ common variants (**Fig. 5e**). We further compared pr_7 genes with biological pathways that are significantly associated with SCZ. We found after removing all the genes that are associated with SCZ pathways, the pr_7 specific genes are still significantly associated with SCZ common variants, and exhibit a similar enrichment level compared to the SCZ pathways specific genes (**Fig. 6d**), demonstrating the enrichment is independent of the annotated SCZ pathways. Furthermore, the pr_7 specific genes include three of the top four significant SCZ rare variant genes (*SETD1A*, *CUL1*, *TRIO*), which are also associated with regulatory function. In contrast to the SCZ pathways genes, pr_7 were also strongly enriched for previously fine mapped genes (**Fig. 6c**). We also utilized a machine learning scheme to nominate the candidate causal variants and found all the identified promoter-associated variants are within pr_7 modules, highlighting the potential causal role of such genes.

In addition to the protein-coding genes, pr_7 contains 1,294 ncRNA, including *DGCR5*, *NEAT1*, *MALAT1*, *MEG3*, and *HAR1A* that have been implicated in neuropsychiatric disorders^51^. The promoter of the ncRNAs exhibits similar enrichment compared to the protein-coding genes for ASD, BD, and SCZ common variants. In addition, we have performed ncRNA-specific MAGMA analysis and found pr_7 ncRNAs are also strongly enriched for ASD and SCZ common variants, suggesting the potential role of such ncRNA in neuropsychiatric disorder. Together, our analysis expanded the genetically susceptible gene set for neuropsychiatric disorders.

Recently, the omnigenic model was proposed to explain the polygenicity in complex traits^58,59^. The model suggests that regulatory genes (peripheral genes) exert their effects by interacting with core genes (e.g. synaptic genes, in the case of neuropsychiatric traits). However, in the case of SCZ, the most significant rare coding variants are associated with regulatory functions, and the fine-mapped gene sets are not restricted to known synaptic pathways. The extent to which the model applies is still highly debatable^60–62^. With chromatin co-accessibility analysis, we identified a chromatin co-accessibility module that is enriched for loss-of-function mutations, rare variants, and previously fine-mapped genes (**Fig. 4**), highlighting the potential causal function for regulatory genes in SCZ. In addition, this module contains genes with similar enrichment for neuropsychiatric disorders (ASD, BD, and SCZ) compared to annotated SCZ pathways. Interestingly, a recent rare variants analysis validated our findings by highlighting RNA processing, chromatin modification, and vesicle-mediated transport as critical pathways for ASD^63^. As such pathways themselves are not significantly associated with risk variants, a plausible explanation is that a relatively limited fraction of genes within such pathways, which have brain-region and cell-type specific chromatin accessibility, contribute to neuropsychiatric disorder heritability.

Overall, our findings highlight the importance of studying CRE alterations across brain regions. As neuropsychiatric traits are closely related to neurodevelopment^64^, and the sorted cells also consist of multiple cell subpopulations, the further integration of single-cell ATAC-seq profiling with different developmental-stage, brain regions, and disease statuses will promise additional insight into the molecular mechanism in neuropsychiatric disorder.

## Acknowledgments

We thank the computational resources and staff expertise provided by the Scientific Computing of the Icahn School of Medicine at Mount Sinai. This study was supported by the National Institute of Mental Health: NIH grants nos. R01-MH109677 (P.R.), U01-MH116442 (P.R. and V.H.), R01-MH125246 (P.R.) and RF1-MH128970 (P.R.), and the National Institute on Aging: NIH grants nos. R01-AG050986 (P.R.), R01-AG067025 (P.R. and V.H.) and R01-AG065582 (P.R. and V.H.). P.D. was supported in part by NARSAD Young Investigator Grant 29683 from the Brain & Behavior Research Foundation. G.E.H. was supported in part by NARSAD Young Investigator Grant 26313 from the Brain & Behavior Research Foundation. J.B. was supported in part by Alzheimer’s Association Research Fellowship AARF-21-722200.

## Author Contributions

P.R. conceived of and designed the project. J.F.F. and P.R. designed experimental strategies for epigenome profiling of human postmortem tissue. D.A.D, V.H., and W.K.S. dissected and provided brain specimens. R.M. prepared nuclei and performed FANS. R.M., Z.S. and J.E. generated ATAC-seq and RNA-seq libraries. S.A. and N.L. performed sequencing of ATAC-seq and RNA-seq libraries. P.D. and P.R. designed analytical strategies. J.B. and P.D. conducted initial bioinformatics, sample processing and quality control. P.D. developed the computational scheme and performed the downstream analysis. J.F.F and P.R. supervised data generation. G.E.H. and P.R. supervised data analysis. P.D. and P.R. wrote the manuscript with input from all authors.

## Supplementary Figures

**Fig S1.**
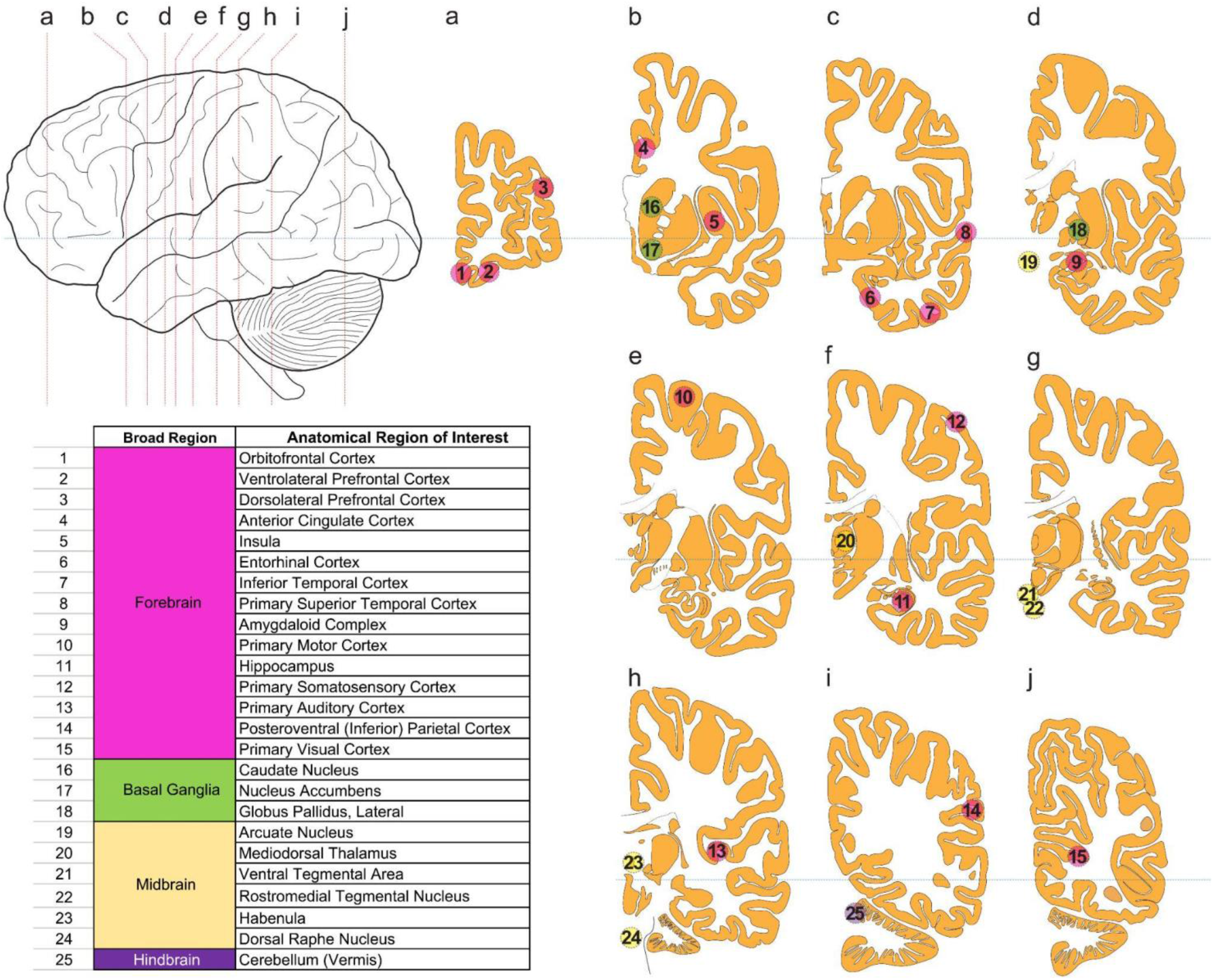
Anatomical localization of the brain regions assayed in the study. Tissue samples used in this study were isolated from 25 functionally distinct brain regions from 6 control individuals.

**Fig S2.**
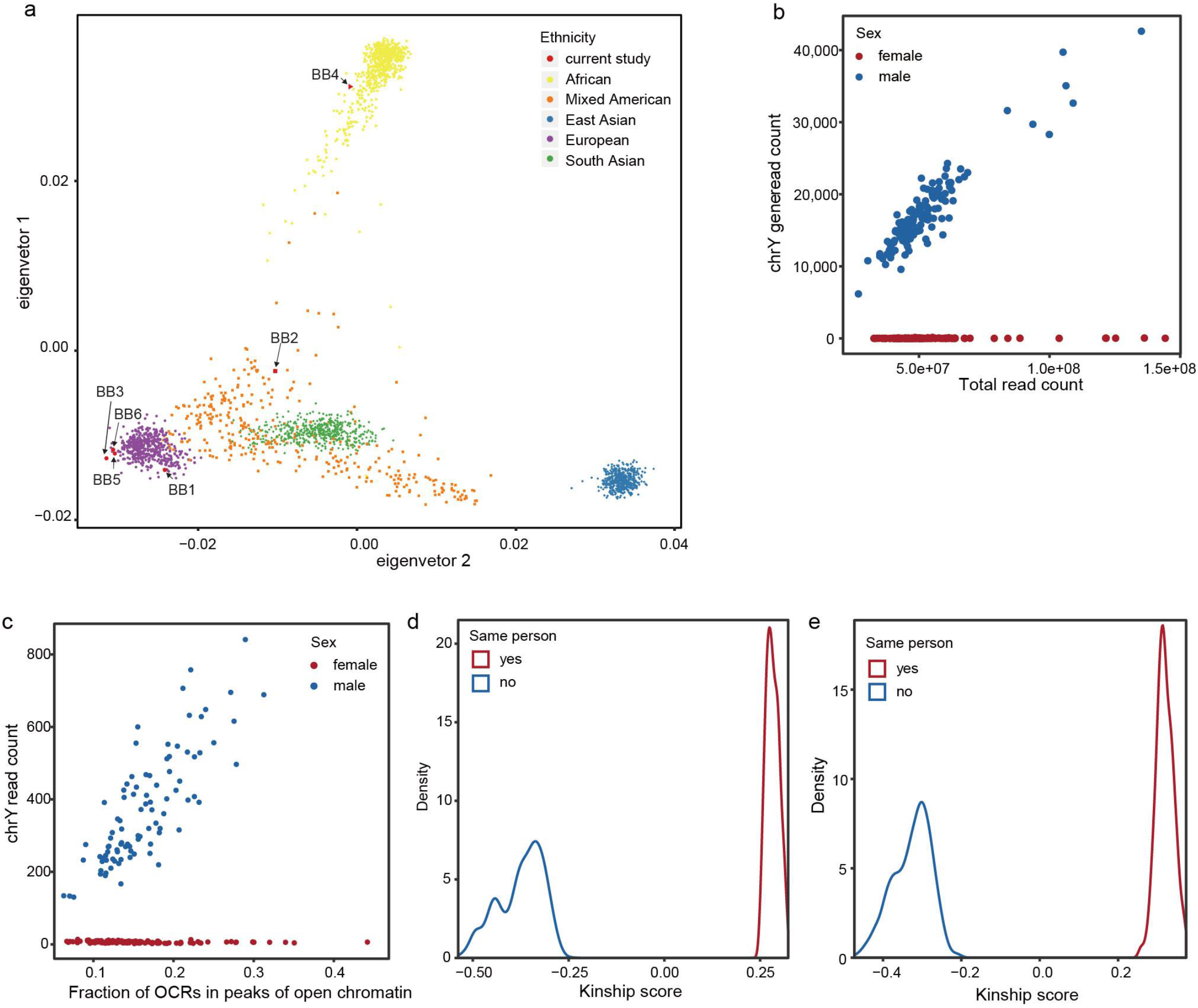
Genetic and sex concordance of the samples. **a**, Principal component analysis of five main 1KG population clusters with individuals from the current study. **b**, RNA-seq sex check based on measuring the number of reads mapped on genes located on chromosome Y (genes located on pseudoautosomal regions are not counted). **c**, ATAC-seq sex check based on measuring the number of reads mapped on chromosome Y. **d**, Genotype concordance (as calculated by KING) based on pair-wise comparison of genotypes called from RNA-seq samples with genotypes from whole-genome sequencing. The histogram shows that none of the non-matching pairs have a positive KING value which indicates full genotype concordance. **e**, Genotype concordance (as calculated by KING) based on pair-wise comparison of genotypes called from ATAC-seq samples with genotypes from whole-genome sequencing. The histogram shows that none of the non-matching pairs have a positive KING value which indicates full genotype concordance.

**Fig S3.**
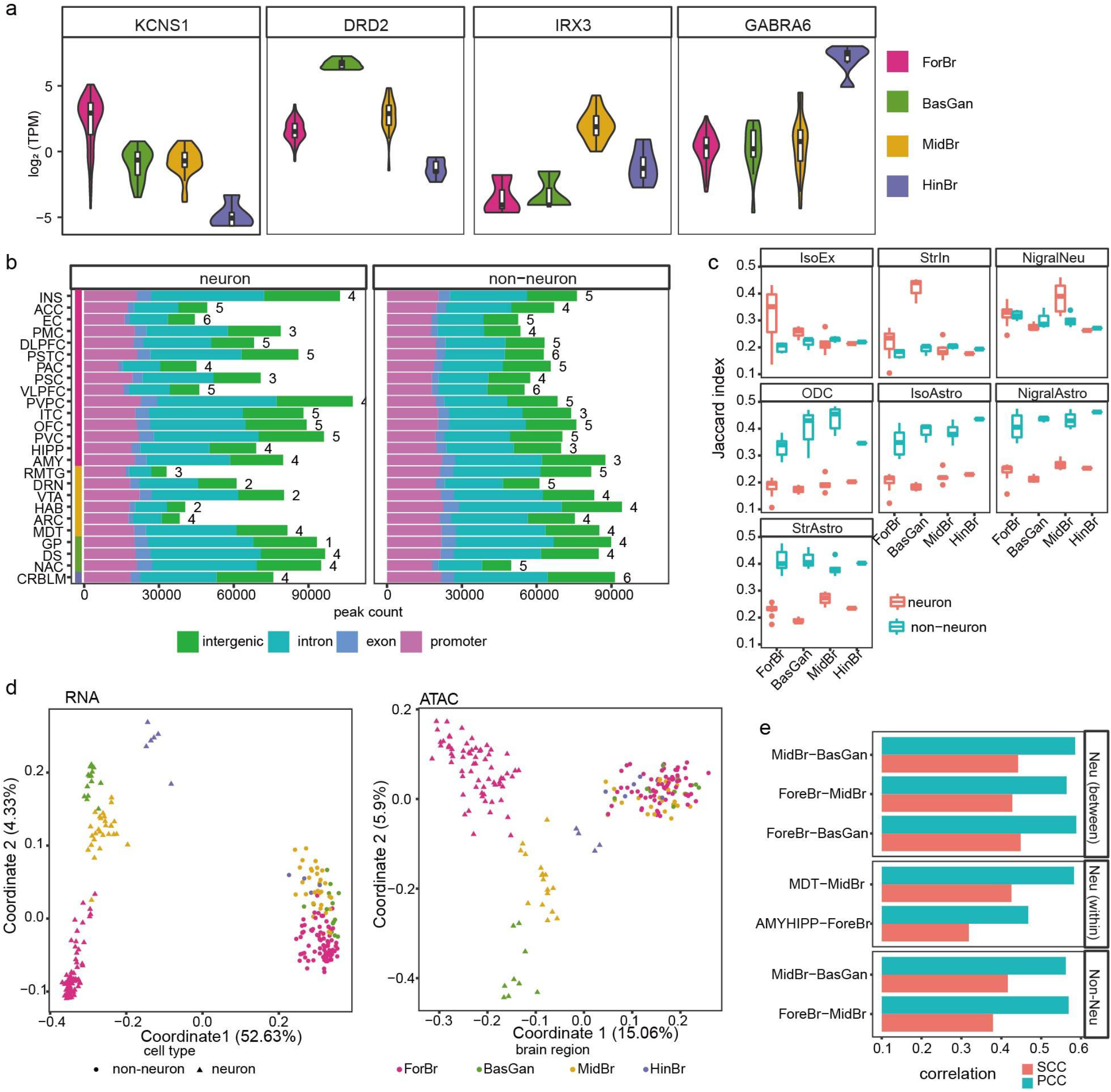
Transcription and chromatin accessibility map across 25 brain regions. **a**, Neuronal gene expression distribution (log_2_TPM) for the brain region-specific marker genes. **b**, Number of peaks and the genomic context profiles merged from the neuronal broad brain regions around (the value right to bar indicates the sample count per group). The promoter is defined as areas within 3kb of TSS. **c**, Jaccard index between our ATAC-seq peaks and a cross-brain region single cell reference^19^, including Isocortical excitatory(ISoEx), Striatal inhibitory (StrIn), Nigral neurons (NigralNeu), ODC (Oligodendrocytes), Isocortical astrocytes (IsoAstro), Nigral astrocytes (NigraLAstro), Striatal astrocytes (StrAstro). **d**, Clustering of the individual samples for ATAC-seq (N=210) and RNA-seq (N=265) using Multidimensional scaling. The value within the prosthesis indicates the percentage of variance explained. **e**, Spearman correlation coefficient (SCC) and Pearson correlation coefficient (PCC) of the log fold change between the gene expression and chromatin accessibility across different comparisons (all P < 2.2 x 10^−16^).

**Fig S4.**
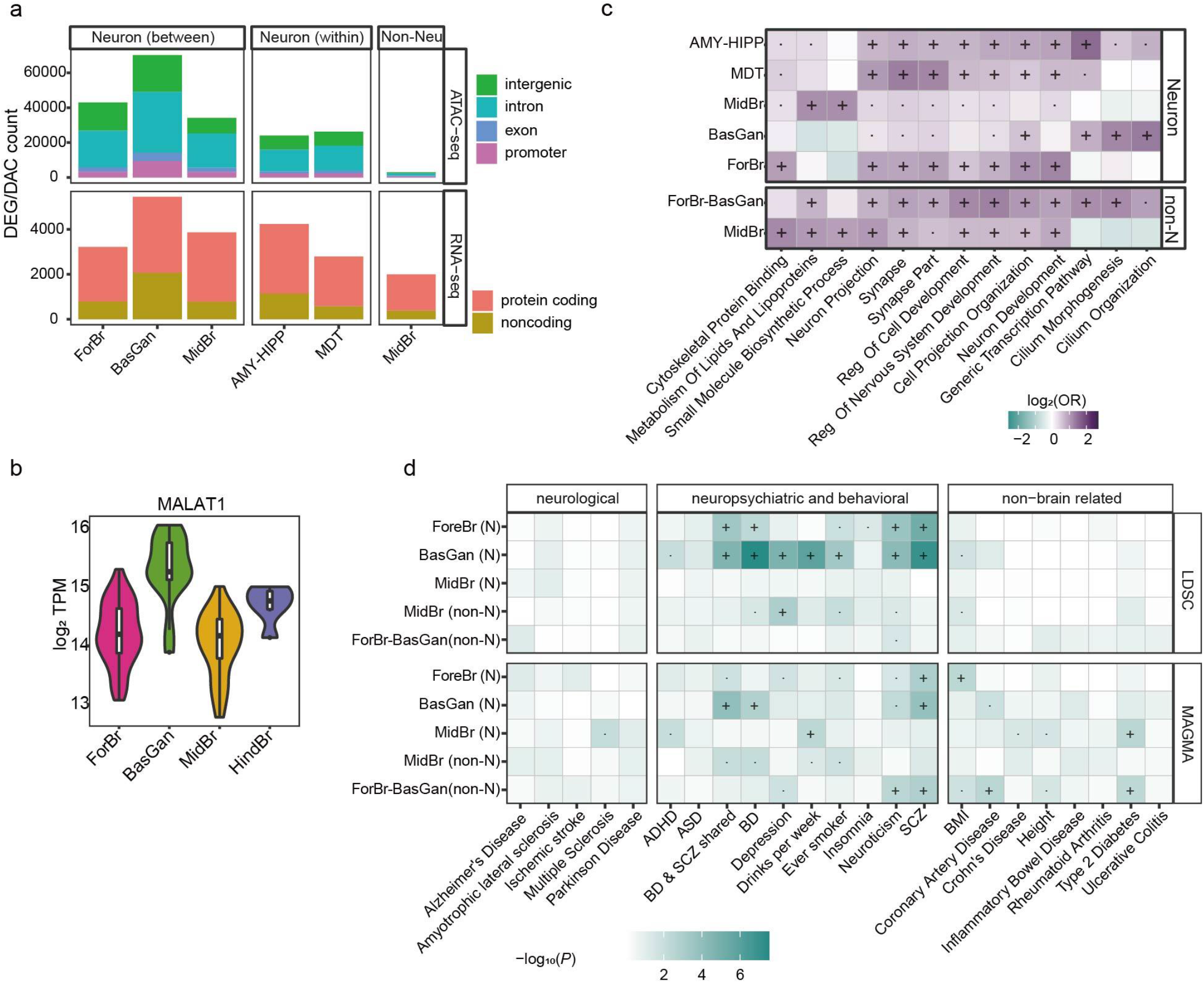
Differential molecular features across brain regions. **a**, Number of differentially expressed genes (DEG) and differentially accessibility chromatin (DAC). For broad brain regions, OCRs/genes were considered specific only if they were significantly more accessible/expressed in all pairwise comparisons against the remaining brain regions. For within broad brain region comparison, only a single pairwise comparison is performed. **b**, gene expression (log_2_TPM) of lincRNA MALAT1 across brain regions neuronal cells. **c**, biological pathway enrichment for the DEGs. non-N represents non-neurons. The color of the heatmap demonstrates the enrichment odds ratio (OR). One-sided fisher-test significance. “·”: Nominally significant (*P* < 0.05); “+”: significant after FDR (Benjamini & Hochberg) correction (FDR < 0.05). non-N denotes non-neurons. **d**, The significance of stratified LD score regression(upper panel) of DAC, and MAGMA (bottom panel) of DEG across different classes of traits for neurons (N) and non-Neurons (non-N).”·”: Nominally significant (*P* < 0.05); “+”: significant after FDR (Benjamini & Hochberg) correction (FDR < 0.05).

**Fig S5.**
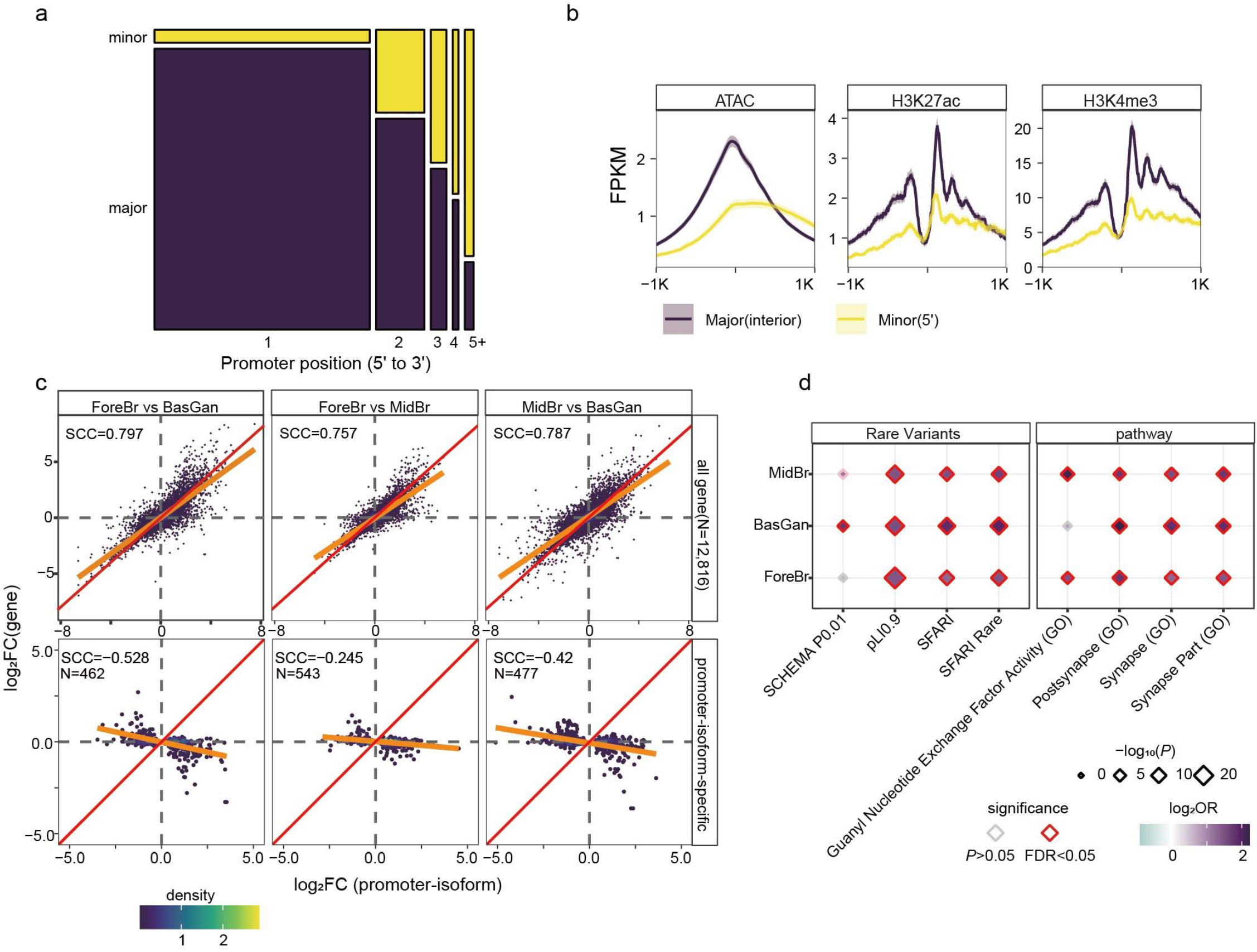
Alternative promoter isoforms. **a**, Major/minor promoter proportions across TSSs ranked by position (5’ to 3’, 1 indicating the most 5’), based on multi-promoter genes with at least one active promoter. **b**, Epigenomic profiles around major interior promoters and non-major 5’ promoters. The shadow shows the 95% confidence intervals.ChIP-seq profiles derived from Dong et al 2022^18^. **c**, Spearman correlation coefficient (SCC) between the log2 fold change of gene and promoter-isoform expressions of all promoter-isoforms (upper panel) and non-concordant promoter-isoforms (bottom panel). **d**, LoF-intolerant genes, rare variants, and biological pathways enriched at non-concordant genes.

**Fig S6.**
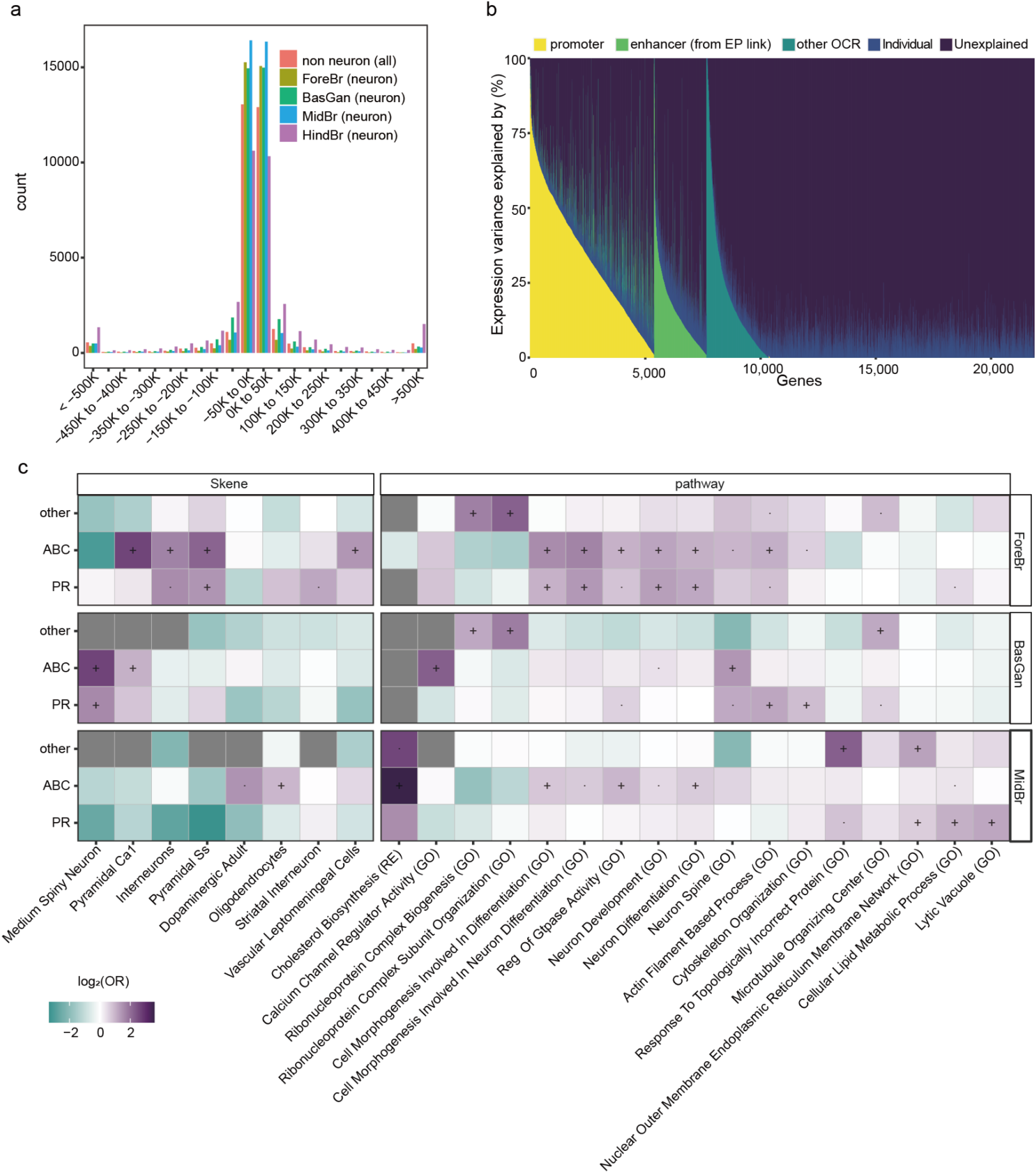
Brain region-specific gene-enhancer links. **a**, Distribution of distance between ABC-enhancer to target genes across different neuronal brain regions, and non-neurons. **b**, Variance component decomposition of the gene expression (every gene per column) by the promoter, enhancer(EP linked), other OCRs within 200kb (other OCR), and individuals of the shuffled samples. This analysis shows that a substantially smaller proportion of variance can be explained by epigenome in the permuted dataset compared Fig. 3c. **c**, Cell-type, and biological pathway enrichment for the brain-region specific DE genes that are explained by the promoter(>%50 variance), EP-linked-enhancer(>%50 variance), and the rest (others). The color of the heatmap demonstrates the enrichment odds ratio (OR). One-sided fisher-test significance. “·”: Nominally significant (p<0.05); “+”: significant after FDR (Benjamini & Hochberg) correction (FDR<0.05).

**Fig S7.**
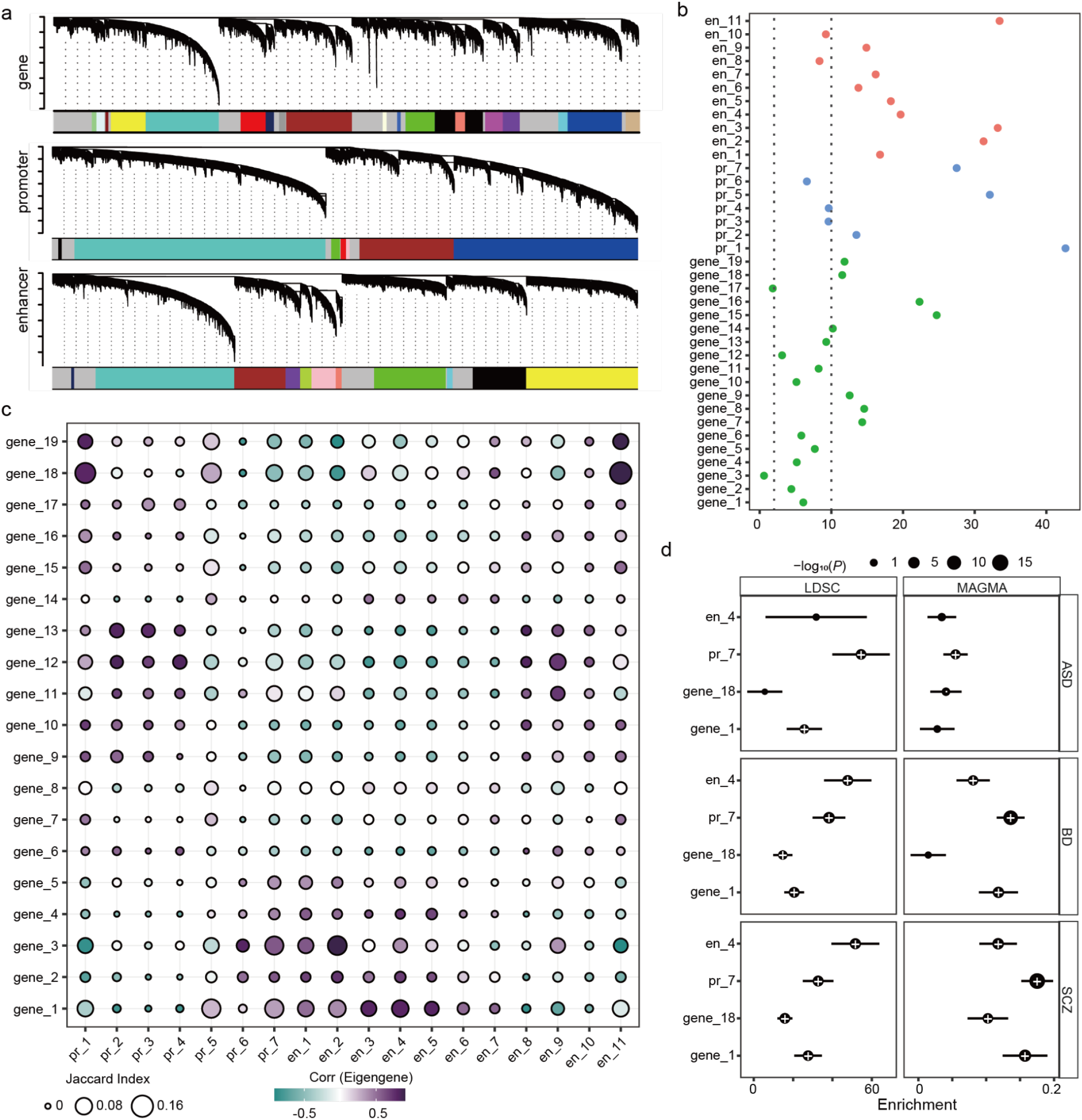
WGCNA. **a**, WGCNA dendrogram for gene, promoter, and enhancer, colored by module. **b**, Z summary score of the preservation of the identified module with independent multi-brain-region RNA-seq and ATAC-seq. (red: promoter, blue: enhancer, green: gene). **c**, The size represents the Jaccard index between genes and promoter/enhancer-linked genes, and the color indicates the correlation of the eigengenes between genes and promoter/enhancer modules. **d**, common variants enrichment of SCZ, ASD, and BD for different categories of genes (determined by MAGMA), CREs (determined by LDSC). A positive coefficient signifies enrichment in heritability (per base enrichment). “·”: Nominally significant (*P* < 0.05); “+”: significant after FDR (Benjamini & Hochberg) correction (FDR<0.05).

**Fig S8.**
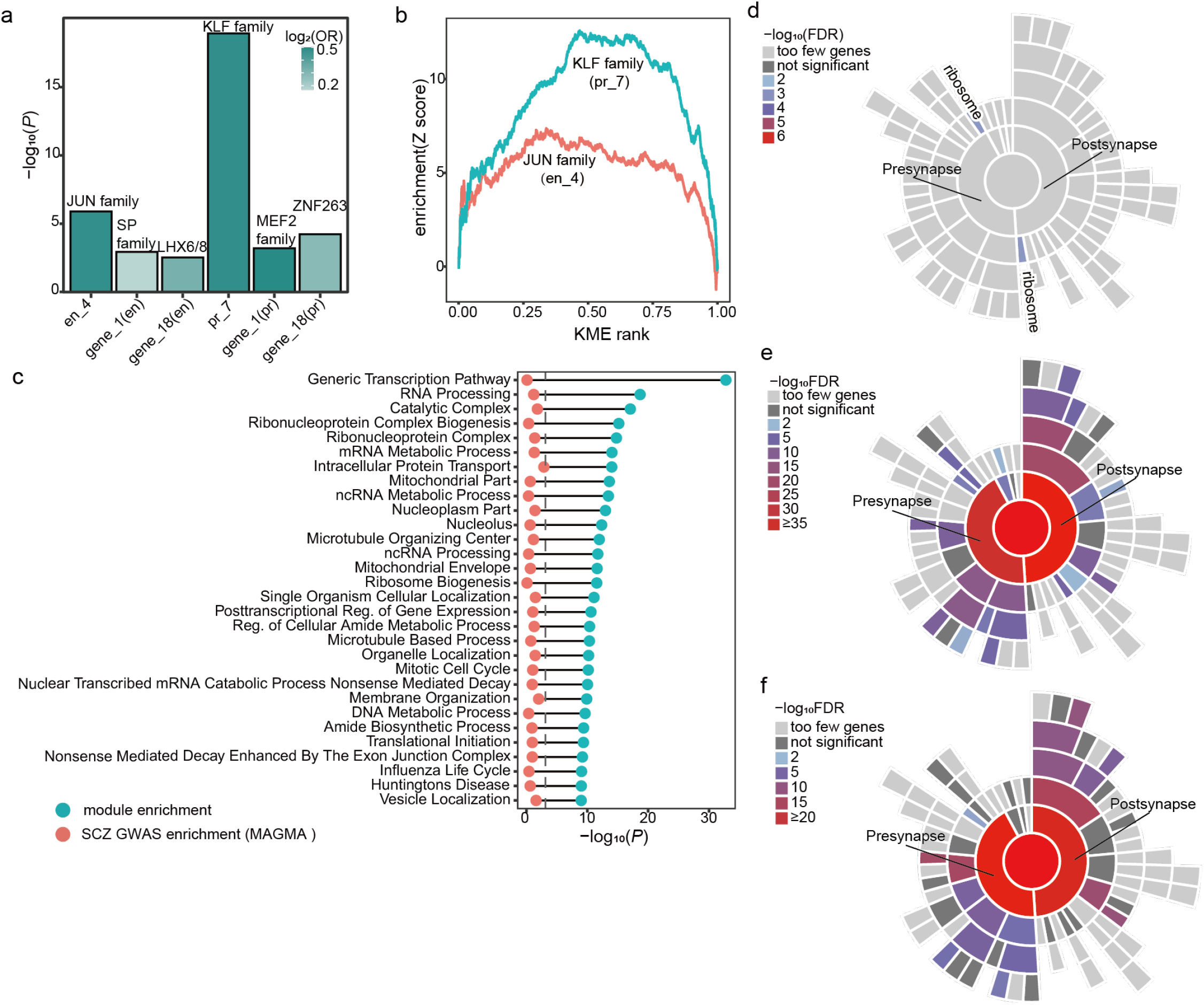
promoter module. **a**, The enrichment (height) and odds ratio (color) of the top enriched TF binding for each of the modules. **b**, Gene-set enrichment analysis plots for the top enriched TF for the CRE module. The extent of deviation from the horizontal reflects the degree of enrichment (+) or depletion (−) at a given centrality (kME). **c**, Top 30 enriched pathways for pr_7, and the pathway enrichment in SCZ common variants (MAGMA, significant after FDR Benjamini & Hochberg correction, FDR<0.05). Sunburst plots depict synaptic locations starting with the synapse (center), pre-and post-synaptic locations in the first ring, and child terms in subsequent rings for **d**, pr7 specific **e**, shared, and **f**, SCZ-pathway-specific genes. Generated by SynGO^65^.

## Methods

### Description of the post-mortem brain samples

Brain tissue specimens from 25 brain regions (**Fig. S1**) were obtained from 4 Caucasian, 1 Hispanic, and 1 Afro-American subjects (determined by self-report and ancestry informative marker analysis) with no history of psychiatric disorder, including alcohol or illicit substance abuse (negative toxicology) or were taking neuropsychiatric medications, including benzodiazepines, anticonvulsants, antipsychotics (typical or atypical), antidepressants or lithium. Four subjects (2x male, 2x female) were collected at autopsy at the Brain Endowment Bank (BEB) at Miller School of Medicine at the University of Miami, and 2 subjects (1x male, 1x female) were collected from Mount Sinai Brain Bank (MSBB) (**Supplementary Table 1**). The cause of death, i.e., sudden cardiac death for all 6 subjects, as determined by the forensic pathologist performing the autopsy. All brain specimens were obtained through informed consent and/or brain donation programs at the Miller School of Medicine at the University of Miami and the Icahn School of Medicine at Mount Sinai. All procedures and research protocols were approved by Institutional Review Boards.

### FANS sorting of neuronal and non-neuronal nuclei

Frozen brain samples were homogenized in cold lysis buffer (0.32M Sucrose, 5 mM CaCl_2_, 3 mM Mg(Ace)_2_, 0.1 mM, EDTA, 10mM Tris-HCl, pH8, 1 mM DTT, 0.1% Triton X-100) and filtered through a 40μm cell strainer. Filtrates were underlaid with sucrose solution (1.8 M Sucrose, 3 mM Mg(Ace)_2_, 1 mM DTT, 10 mM Tris-HCl, pH8) and subjected to ultracentrifugation at 24,000 rpm for 1 hour at 4°C. Pellets were resuspended in 500μl DPBS and incubated in BSA (final concentration 0.1%) and anti-NeuN antibody (1:1000, Alexa488 conjugated, Millipore, MAB377X) under rotation for 1 hour, at 4 °C. Just prior to FACS sorting, DAPI (Thermo Scientific) was added to a final concentration of 1μg/ml. Neuronal (NeuN+) and non-neuronal (NeuN-) nuclei were sorted using FACSAria (BD Biosciences).

### Generation of RNA-seq libraries

RNA was isolated from 149 tissue dissections from 25 brain regions. RNA-seq libraries were generated using the SMARTer Stranded Total RNA-seq kit v2 (Takara Cat no. 634419) according to the manufacturer’s instructions. Libraries were sequenced with the Illumina Hiseq using 100bp paired-end reads. After quality controls (see below), we retained 265 RNA-seq libraries, which, on average, corresponds to available RNA-seq data for 22 out of the 25 regions per individual.

### Processing of RNA-seq libraries

Each set of pair-end reads was processed by Trimmomatic^66^ to remove low-quality base pairs and sequence adapters. Reads were subsequently aligned to the human reference genome GRCh38 using STAR^67^. To correct for allelic bias resulting from individual-specific genome variation, we ran STAR with the enabled WASP module^68^ as we provide both RNA-seq FASTQ file and the Whole Genome sequencing (WGS) file of the corresponding individual. The BAM files that were generated contain the mapped paired-end reads, including those spanning splice junctions. Following read alignment, expression quantification was performed at the transcript isoform level using RSEM^69^ and then summarized at the gene level. Gene quantifications correspond to GENCODE^70^. Quality control metrics (**Supplementary Table 3**) were reported with RNA-SeqQC^71^, Qualimap^72^, and Picard.

### Generation of ATAC-seq libraries

ATAC-seq reactions were performed on Neuronal (NeuN+) and non-neuronal (NeuN-) nuclei isolated by FANS from 149 tissue dissections from 25 brain regions, resulting in 291 ATAC-seq libraries. Where available, 100,000 sorted nuclei were centrifuged at 500 × g for 10 min, 4°C. Pellets were resuspended in the transposase reaction mix and libraries were generated using an established protocol^73^. Libraries were sequenced with the Illumina Hiseq 4000 using 50bp paired-end reads. After quality controls, we retained 210 ATAC-seq libraries, which, on average, corresponds to available ATAC-seq data for about 18 out of the 25 regions per individual.

### Processing of ATAC-seq libraries

The raw reads were trimmed with Trimmomatic^66^ and then mapped to the human reference genome GRCh38 analysis set reference genome with the pseudoautosomal region masked on chromosome Y with the STAR aligner^67^. To correct for allelic bias resulting from individual-specific genome variation, we ran STAR with enabled WASP module^68^ as we provide both ATAC-seq FASTQ file and WGS file of the corresponding individual. This yielded for each sample a BAM file of mapped paired-end reads sorted by genomic coordinates. From these files, reads that mapped to multiple loci or to the mitochondrial genome were removed using samtools^74^ and duplicated reads were removed with PICARD. Quality control metrics (**Supplementary Table 4**) were reported with phantompeakqualtools^75^ and Picard.

### Analysis of differentially expressed/accessible genes, promoter-isoforms, and OCRs

To assess which genes, isoforms, and OCRs showed differential expression and accessibility, we employed statistical modeling based on linear mixed models. The starting point here was three count matrices with raw read counts per each sample and feature (i.e., gene, promoter-isoform, or OCR). For gene expression and chromatin accessibility, we excluded features that were lowly expressed/accessible by only keeping those with at least 1 count per million(CPM) reads in at least 20% of the samples. For promoter-isoforms, we used a more stringent threshold, and only kept those with at least 2 CPM reads in >40% of the samples. Then, the read counts were normalized using the trimmed mean of M-values (TMM) method^76^. Covariates were selected as implemented in our previous study^9^ (**Supplementary Table 5**). Statistical Analysis of differences in gene/isoform expression and chromatin accessibility: The normalized read count matrices from voomWithDreamWeights (variancePartition package^78^) was then modeled by fitting weighted least-squares linear regression models estimating the effect of the right-hand side variables on the expression/accessibility of each feature.

As our dataset contains up to 25 samples per individual within both neuronal and non-neuronal subsets, we ran differential analysis by dream^77^ method from variancePartition package^78^. Dream properly models correlation structure and, thus, keeps the false discovery rate lower than the other commonly used methods for this purpose. Finally, adjusted matrices of gene/promoter-isoform expression and chromatin accessibility were created for neuronal and non-neuronal samples where the effects of Sex and technical covariates were removed.

### Canonical Correlation Analysis (CCA)

CCA between gene expression and chromatin accessibility was performed based on R toolkit Seurat^79^ and Signac^80^. Briefly, we determined gene activity score for all the expressed genes from chromatin accessibility across each sample. We focused on the protein-coding genes that are highly variable for gene expression (top 2000 dispersion, aka variance to mean ratio) or gene activity (top 2000 dispersion). Then we utilized a variant of CCA, diagonal CCA^29^ to construct our canonical correlation vectors.

### Promoter-isoform analysis

We employed the proActiv package to determine the promoter-isoform expression^32^. Briefly, we first identified the uniquely identified promoter-isoform (**Supplementary Table 6**), which excludes the isoform that is a single-exon or uses an internal intron (first intron edge of current isoform overlapped with non-first intron edge of other isoforms). We assign the isoform-transcript id as the promoter-isoform; when a promoter-isoform corresponds to multiple isoforms, we randomly choose one. Then we quantified the promoter-isform expression by estimating the junction reads aligned to the first intron. We used the same differential analysis for promoter-isoform analysis. To determine the promoter-isoform-specific DEGs, we focused on the significant (FDR <0.05) promoter-isoforms between broad region pairwise comparisons. We selected such promoter-isoforms that the genes are not significant (nominal *P* > 0.1) or have an opposite fold change direction (**Supplementary Table 7**).

### Activity-by-contact (ABC) gene-enhancer links

To determine the target gene for a distal CRE, we utilized the Activity-by-contact model^35^ which integrates chromatin states and chromosome spatial organization information to construct gene-enhancer links for ForeBr (neuron), MidBr (neuron), BasGan (neuron), HindBr (neuron), and non-neuron (merged from all three brain regions) cells. If the unique identified promoter-isoform annotation is available, we utilized the most highly expressed promoter-isoform to define the promoter for the gene. As cell-type confers larger variance compared to brain regions, we utilized the Hi-C of neuronal and non-neuronal cells as input^18,36^. In accordance with the authors’ directions, we filtered out predictions for genes on chromosome Y and lowly expressed genes (genes that did not meet inclusion criteria in our RNA-seq dataset). We used the default threshold of ABC score (a minimum score of 0.02) and the default screening window (5MB around the TSS of each gene).

As the model only considered spatial proximity and epigenomic signal strength, we next assessed the enhancer-gene coordination. We first compared the correlation between gene expression and chromatin accessibility for brain region-specific EPs, brain-region-shared EPs, and non-EPs for each brain region. We found that these correlations are much higher for both brain region-specific-EPs and shared EPs compared to the background (non-EPs).

### Variance explain

We used variance component analysis to assess the extent to which gene expression variability could be explained by promoter, ABC enhancer, and other OCRs following an implementation suggested by a previous report^37^. A variance component analysis was used to examine how much gene expression variability could be correlated to patterns of chromatin covariance (Fig. 2). To implement such a model, we followed an implementation suggested by a previous report^37^. First, we modeled negative binomial distribution of RNA-seq count data by a variance stabilizing transformation (vst; varistran R package (v.1.0.4)). Then, for each gene represented by vst-normalized vector g, we considered the following variance component model: 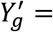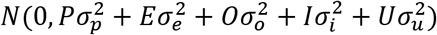

where P, E, and O are sample-sample covariance matrices of chromatin accessibility in promoter (OCRs overlapping region within 1kb from the transcription start site), ABC enhancers, and other distal OCRs (not promoter or ABC enhancer, and overlapped within 1-100kb from the transcription start site), 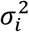 captures per-individual covariance matrix and 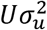 is the noise term. The values of 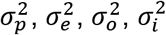, and 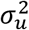, were estimated by the average information restricted likelihood estimation (AIREML; gaston R package (v.1.5.5)). We used residualized count matrices of OCRs where the effect of technical covariates was regressed out. For clarification, this approach does not model the relationship of each gene to its own promoter/enhancer OCRs but to the overall status of all enhancers/promoter OCRs.

### Partitioned heritability with stratified LD score regression

We partitioned heritability for DE peaks/TEns as well as top eSNPs to examine the enrichment of common variants in neuropsychiatric traits with stratified LD score regression (v.1.0.1)^44^ from a selection of GWAS studies, including:

Attention-deficit/hyperactivity disorder (ADHD)^81^, Autism spectrum disorder(ASD)^3^, Bipolar Disorder and Schizophrenia shared (BD & SCZ shared)^82^, Bipolar Disorder (BD)^2^, Major depression^83^, Drink per week^84^, Ever smoker^84^, Insomnia^85^, neuroticism^86^ Schizophrenia (SCZ; EAS ancestry, EUR ancestry, and merged)^1^, Alzheimer’s Disease^87^, Amyotrophic lateral sclerosis^88^, Multiple Sclerosis^89^, Parkinson Disease^90^, body mass index (BMI)^91^, Coronary Artery Disease^92^, Crohn’s Disease^93^, Height^94^, Inflammatory Bowel Disease^93^, Ischemic stroke^95^, Rheumatoid Arthritis^96^, Type 2 Diabetes^97^, Ulcerative Colitis^93^.

Briefly, with the input peaks, a binary annotation was created by marking all HapMap3 SNPs^98^ that fell within the peak or eSNPs and outside the MHC regions. LD scores were calculated for the overlapped SNPs using a LD window of 1cM using 1000 Genomes European Phase LD reference panel^99^. The enrichment was determined against the baseline model^44^. We also determined the gene sets enrichment with the same pipeline (**Fig. 6d**). The genes were padded by 35kb upstream and 10kb downstream, and HapMap3 SNPs that fell within such regions and outside of the MHC regions were utilized. To compare the GWAS enrichment between populations, we obtained the EUR and EAS SCZ GWAS summary statistics^1^ and determined the enrichment with ancestry-specific LD reference and baseline model provided by the software website.

### Overlap of OCR with existing annotation

We collected cell-type-specific cross-brain region OCRs from a recent single-cell ATAC-seq reference^19^. First, we compared the Jaccard index between our OCRs with the reference and confirmed the cell-type and brain region specificity. In addition, we utilized the union of all peaks as background and performed a single side fisher exact test between the DAC and the single-cell makers.

### Gene set enrichment analysis

To explore the function of a gene set, we collected functional gene sets from i) MSigDB 7.0^100^, ii) human brain single-cell markers^21,50^(For Skene et al, top 1 specificity percentile were used), SCZ common variants colocalized or fine mapped gene sets (PGC3 fine map^1^, PEC high confidence^7^, PEC TWAS^48^, fetal brain fine mapped TWAS ^47^), interoperate for loss-of-mutation (defined as pLI > 0.9)^49^, TF genes (GO term GO:0003700, GO:0004677, GO:0030528, GO:0004677, GO:0045449), and rare variants (SCHEMA^46^ P < 0.01, SAFARI). One-tailed Fisher exact tests were used to test the enrichment and significance. and synaptic gene ontology resource^65^. In addition, we have performed the gene set enrichment with SynGO^65^ with default parameters.

To examine the genetic enrichment of gene sets, we used MAGMA (v 1.07b)^45^ with GWAS data (described above). Briefly, genes were padded by 35kb upstream and 10kb downstream, and the MHC region was removed due to its extensive linkage disequilibrium and complex haplotypes. The European panels from 1000 Genome Project phase 3 were used to estimate the Linkage disequilibrium (LD)^99^. The BETA value from the MAGMA output was used to represent the enrichment. To determine the pathways that are enriched for SCZ common variants, we determined the MAGMA enrichment for all the above-mentioned MSigDB pathways and collected all the significant pathways after FDR (Benjamini & Hochberg) correction (FDR < 0.05)^101^. In parallel, we have also determined the ncRNA enrichment with the same pipeline, independent of the gene analysis^102^. Only the brain-expressed ncRNAs (as determined by the FANTOM project) were utilized in this analysis. We determined the EUR and EAS SCZ GWAS enrichment with ancestry-specific GWAS summary statistics and sub-poupulation 1,000 Genomes references. For the conditional analysis, we used the condition modifier of the ‘gene-covar’ parameter to condition on the normalized gene expression of the reported cell types, including hippocampal CA1 pyramidal cells, striatal MSN, pyramidal SS, and cortical interneurons^1,21^.

### WGCNA network construction

We constructed a gene co-expression and chromatin co-accessbility network using Weighted Gene Co-expression Network Analysis (WGCNA)^103^. All covariates, except for brain region, were regressed out of the gene expression and chromatin accessibility matrix (log2 CPM). For scalability, we construct enhancer and promoter networks independently. Briefly, the power parameters were determinedas the smallest power(from 5 to 20) that achieves a truncated R^2^ of 0.8. The network dendrogram was created using average linkage hierarchical clustering of the topological overlap dissimilarity matrix (1-TOM) with hybrid dynamic tree-cutting. Modules were defined using biweight midcorrelation (bicor), with a minimum module size of 100, deepsplit of 2, merge threshold of 0.1. The initial modules were then merged based on the eigengene with the “ mergeCloseModules” function. Module membership (kME) were determined by the correlation to the module eigengene (**Supplementary Table 10 - 12**). The top ranked genes or OCR-linked genes (‘hub’ genes) for the selected modules were shown in Fig. 4b. As an OCR can be linked to multiple genes, all the genes were shown.

To assess the module preservation, we examined the Z-summary statistics^104^ with independent multi-brain region RNA-seq from GTEx^105^ (ForeBr: Cortex, Amygdala, Hippocampus, Frontal Cortex, Anterior cingulate cortex; BasGan: Caudate, Putamen, Substantia nigra; MidBr: Hypothalamus) and ATAC-seq^9^ (ForeBr: ACC, DLPFC, INS, ITC, OFC, PMC, VLPFC, PVC,AMY, HIPP; BasGan, Nac; MidBr: MDT). It’s worth noting the GTEx RNA-seq data contains both neuron and non-neuron cells. A Z summary score < 2 indicates the lack of preservation.

### Module membership-based gene set enrichment

To determine the association between module membership and the enrichment of Loss-of-function intolerant genes and TF genes, we utilized a Brownian-Bridge-statistics-based method ^62^. Briefly, the genes (or OCR-linked genes) were ranked by the membership (the OCR’s membership in the case of OCR-linked genes), and the proportions of the genes that are within the set (LOF or TF, r) is determined. At a given quantile q of genes, we then determined how many of the genes are within the set Obs(q). The Expected number of genes within a set would be E(q)=q*r*M, and variance V(q)=q*(1-q)*M*r*(1-r). The *Z*-score can be determined subsequently as 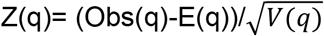. We converted the Z-scores to *P* values, and the significance is determined with the minimum *P* value after FDR (Benjamini & Hochberg) correction (FDR < 0.05)

### gkm-SVM variant effect identification

We combined gkm-SVM^52^ with three complementary approaches, GkmExplain^53^, in silico mutagenesis^54^, and deltaSVM^55^ following a recent analysis ^19^. Briefly, we used run gkm-SVM^52^ model with a 10 fold cross-validation on 1000bp loci centered around ATAC-seq summits and an equal size GC content matched non-OCRs for each neuronal brain region to classify the sequences that are accessible or not. Then, we collected the SNPs that are genome-wide significant (P < 5 × 10^−8^) or fine-mapped causal variants (posterior probability > 0.01) from the most recent SCZ GWAS result^1^, as well as any SNPs in linkage disequilibrium (LD) with the 2 categories (defined as LD R^2^ > 0.8). Lastly, we determined the score from GkmExplain^53^, in silico mutagenesis^54^, and deltaSVM^55^, and selected the SNPs that are significant with all three methods.

## Data availability

The datasets generated in the current study are available through the Gene Expression Omnibus (GEO) under accession number GSE211826 (ATAC-seq and RNA-seq), and Sequence Read Archive (SRA) under accession number PRJNA870417 (WGS. The UCSC genome browser tracks of our processed ATAC-seq data and download links are available at our webpage (https://dongp01.u.hpc.mssm.edu/multiregion.html).

The following publicly available datasets were used: human multi-brain-region single-cell ATAC-seq reference^19^ (NCBI GEO GSE147672), mouse multi-brain-region single-cell RNA-seq reference^21^, human brain single-cell RNA-seq reference^50^. Human multi-brain region RNA-seq data from GTEx^105^ (https://gtexportal.org/home/), human multi-brain region ATAC-seq data^9^ (NCBI GEO GSE96949).

