## Supplementary Notes for "Transcriptome and chromatin accessibility landscapes across 25 distinct human brain regions expand the susceptibility gene set for neuropsychiatric disorders"

### Supplementary Methods

##### Generation of whole-genome sequencing libraries

Whole-genome sequencing libraries were prepared at BGI Genomics. DNA was extracted from frozen tissue sections using the QIAamp DNA mini kit (Qiagen, Cat no. 51306) according to the manufacturer's instructions. Whole-genome data were generated for all 6 subjects on Illumina Hiseq 4000 using 100bp paired-end reads.

##### Data processing and quality control of whole-genome sequencing libraries

To facilitate the alignment of raw sequencing files and perform variant calling, we utilized CCGD pipeline (<https://github.com/CCDG/Pipeline-Standardization/>)[^1^](https://sciwheel.com/work/citation?ids=5930905&pre=&suf=&sa=0). In brief, reads were aligned to hg38 human reference genome using BWA-MEM. Then, the pipeline follows GATK Best Practises guidelines to perform duplicate marking, base recalibration, indel-realignment, quality score binning, and variant calling. All samples were noted to have broadly even profiles across quality control metrics, i.e genome coverage > 98% (10x coverage > 96% loci), dbSNP coverage > 98%, and transition/transversion (Ti / Tv) ratios between 2.07 and 2.08, consistently with our genome-wide expectations (general range for known and novel loci on the human genome is ∼2.0-2.1[^2^](https://sciwheel.com/work/citation?ids=148564&pre=&suf=&sa=0), while the empirical value for Illumina Hiseq platform is 2.07[^3^](https://sciwheel.com/work/citation?ids=1820479&pre=&suf=&sa=0)) (**Supplementary Table 2**).

To verify ancestry information, we merged the whole genome samples with 1KG cohort (http://ftp.1000genomes.ebi.ac.uk/vol1/ftp/release/20130502/supporting/GRCh38_positions/) and performed principal component analysis on the thinned set of 30,000 randomly selected SNPs with MAF ≥ 5% (SNPRelate package v1.16[^4^](https://sciwheel.com/work/citation?ids=1038588&pre=&suf=&sa=0)). Based on the proximity of our samples to 1KG population clusters in the two-dimensional space of the first two principal components, we checked the ethnicity information of all six individuals (**Fig. S2a**).

##### Quality control of RNA-seq libraries

On average, over 51 million sequenced paired-end reads were obtained for each sample. In our initial dataset of 308 samples, 17 samples had a technical or biological replicate. We decided to keep those replicates with a better correlation to the gene expression profiles of the other samples originating from the same cell type and brain region. In case of similar results, we retained a sample with a higher number of uniquely mapped reads. Then, we calculated the correlation of each sample to all other samples of the same cell type and highlighted 26 samples with markedly different correlations, i.e., the difference in mean correlation of the given sample with the rest of the dataset and the mean correlation of all pairs of samples within the dataset was more than twice higher than the standard deviation calculated upon all pair’s correlation. All those 26 samples originating from 15 distinct brain regions of 5 individuals were removed after a visual inspection in IGV combined with an inspection of metadata which identified probable reasons such as the low amount of starting material, low RIN, or low ratio of uniquely mapped reads. When we applied this filtering procedure, we ended up with a final set of 265 samples, i.e., 132 neuronal and 133 non-neuronal (**Supplementary Table 3**) that are well separated in PCA (**Fig. S3d**). To check the sex of individuals within our cohort, we also measured the number of reads mapped on genes located on chromosome Y (genes located on pseudoautosomal regions are not counted; **Fig. S2b**). To check the identity of RNA-seq samples, we ran genotype comparisons of all samples against each other and against imputed genotypes from whole genome sequencing (WGS) (**Fig. S2d**).

##### Quality control of ATAC-seq libraries

On average, over 57 million sequenced paired-end reads were obtained for each sample. Because of using FANS sorted nuclei as opposed to whole cells, only a low fraction of the reads were mapped to the mitochondrial genome (mean of 0.97% of the uniquely mapped reads). In our initial dataset of 293 libraries, seven libraries had a technical or biological replicate. We decided to keep those replicates with a better correlation to the chromatin accessibility profiles of the other samples originating from the same cell type and brain region. We also excluded libraries that had low mappability (less than 50%), low per-sample called OCRs (less than 3,000), low GC content (less than 90% of cell type median, i.e., 52.15% and 54.35% for neuronal and non-neuronal libraries, respectively) or low final read count (less than 5,000,000). The threshold of cell GC content was set empirically by testing all values between 75-95% of cell-type median (with a step of 5%) and observing changes in the clustering analysis as those samples with low GC were frequently outliers in the MDS plots. When we applied this filtering procedure, we ended up with a final set of 210 samples, i.e., 97 neuronal and 113 non-neuronal (**Supplementary Table 4**). Those neuronal and non-neuronal samples are relatively well separated in PCA (**Fig S3d**). Similarly to RNA-seq, we performed a sex check (**Fig. S2c**) and a genotype check **Fig. S2e**). ATAC-seq QC metrics are summarized in Supplementary Table 4.

For further steps, we split samples into neuronal and non-neuronal datasets. The rationale for this decision comes from the differential analysis that was unable to properly correct for the effect of markedly different chromatin compositions of these cell types.

The samples from the same brain region and cell type were subsequently subsampled and merged, creating 50 BAM files with a uniform depth of 170 million paired-end reads. For neuronal samples from globus pallidus (GP), habenula (HAB), ventral tegmental area (VTA), and Dorsal Raphe Nucleus (DRN), we had less than 170 million pair-end reads (39, 123, 136, and 162 million, respectively) so we retained all reads from the corresponding samples. Except for those samples, all subsampling ratios were calculated per each sample individually within those 50 respective groups (brain region by cell type) to ensure that each of them contributes the same number of reads regardless of their overall read counts. Using these BAM files, bigWig files were created, and peaks were called by MACS2[^5^](https://sciwheel.com/work/citation?ids=57981&pre=&suf=&sa=0). After removing peaks overlapping ENCODE blacklisted regions of anomalous, unstructured, or high signal in functional genomics assays[^6^](https://sciwheel.com/work/citation?ids=7132831&pre=&suf=&sa=0), 320,308 and 196,467 peaks remained for neuronal and non-neuronal datasets. For each peak, we assigned the closest gene and the genomic context of an ATAC-seq OCR using ChIPSeeker[^7^](https://sciwheel.com/work/citation?ids=1308084&pre=&suf=&sa=0); the transcript database was built by GenomicFeatures[^8^](https://sciwheel.com/work/citation?ids=790825&pre=&suf=&sa=0) upon ENSEMBL genes. Finally, read counts of all samples were quantified within these peaks using featureCounts function in RSubread[^9^](https://sciwheel.com/work/citation?ids=148598&pre=&suf=&sa=0).

##### Genetic concordance analysis

To verify the identity of samples across all assays, we compared called genotypes of RNA-seq and ATAC-seq samples against whole-genome sequencing samples using KING v1.9[^10^](https://sciwheel.com/work/citation?ids=1433823&pre=&suf=&sa=0). To overcome the issue of a relatively high error rate for variant calling in functional genomics assays, we utilized GATK Best Practises guideline (https://software.broadinstitute.org/gatk/best-practices/workflow?id=11164), followed by the removal of variants with minor allele frequencies (MAF) < 25%. For the RNA-seq cohort, this analysis resulted in the correction of genotypes of 2 unambiguously swapped samples and the removal of 2 samples due to genotype contamination, i.e. high genetic concordance of a single sample with multiple distinct genotypes. In the case of ATAC-seq, we corrected the genotypes of 4 unambiguously swapped samples and removed 5 likely contaminated samples.

##### Covariates selection and differential analysis

The starting point for statistical modeling of gene/isoform expression and chromatin accessibility was chosen with the variables Brain_region (25 levels) and Sex (2 levels) for a base model. Sex was included as it is known to have a strong effect on a few genes/promoter-isoforms/OCRs (features) primarily located on the sex chromosomes. To assess which covariates should be included in order to have a good average model for gene/isoform/OCR accessibility, we employed the Bayesian information criterion (BIC) as implemented in our previous study[^11^](https://sciwheel.com/work/citation?ids=5486493&pre=&suf=&sa=0). This procedure pinpoints the best performing covariates upon the initial sets of 79 and 51 covariates for RNA-seq and ATAC-seq datasets, respectively. We required to net improve at least 5% of the features showed a change of 4 in the BIC score, which is above the lower boundary of “positive” evidence against the null hypothesis. Summaries of selected covariates for all combinations of analysis type and cell type are provided in Supplementary Table 5.
